# Structural insights into transcription activation mechanism of the global regulator GlnR from actinobacteria

**DOI:** 10.1101/2023.01.09.523197

**Authors:** Jing Shi, Zhenzhen Feng, Juncao Xu, Fangfang Li, Yuqiong Zhang, Aijia Wen, Fulin Wang, Qian Song, Lu Wang, Shuang Wang, Yu Feng, Wei Lin

## Abstract

GlnR, an OmpR/PhoB subfamily protein, is an orphan response regulator that globally coordinates the expression of genes responsible for nitrogen, carbon and phosphate metabolism in actinobacteria. Although much efforts at biochemical and genetic analyses have been made on the mechanism of GlnR-dependent transcription activation, it still remains unclear owing to lacking the structure of GlnR-dependent transcription activation complex (GlnR-TAC). Here, we report a crystal structure of a binary complex including a C terminal DNA binding domain of GlnR (GlnR_DBD) and its regulatory *cis*-element DNA, and a cryo-EM structure of GlnR-TAC comprising of *Mycobacterium tuberculosis* RNA polymerase, GlnR, and a promoter containing four well-characterized conserved GlnR binding sites. These structures show four GlnR protomers coordinately engage promoter DNA in a head-to-tail manner, with two N-terminal receiver domains of GlnR (GlnR-RECs) jointly act as a bridge to connect RNAP αNTD with the upstream GlnR_DBD. GlnR-TAC is stabilized by complex protein-protein interactions between GlnR and the conserved β flap, σ^A^R4, αCTD, αNTD domains of RNAP. These are in good agreement with our mutational and kinetic single-molecule fluorescence assays. Altogether, our results reveal a general transcription activation mechanism for the global regulator GlnR and other OmpR/PhoB subfamily proteins, and present a unique mode of bacterial transcription regulation.

**Significance statement:** In low-GC gram-negative bacteria, the typical two component system NtrB/NtrC accounts for the expression of genes related to nitrogen metabolism. In high-GC gram-positive actinobacteria, GlnR, an atypical and orphan response regulator (RR) of the OmpR/PhoB subfamily proteins, globally coordinates transcription of genes involved in nitrogen, carbon and phosphate metabolism. Here, using crystallography, cryo-electron microscopy, and single-molecule fluorescence assays, we show that GlnR activates transcription by contacting DNA between the −10 and −35 elements and further upstream through contacting σ region 4 and RNAP β flap subunit. We also identify a previously unobserved cooperative engagement of four GlnR protomers to the promoter DNA, which not only makes the transcription initiation complex (RPo) more stable, but also provides better transcription activities.

## Introduction

Actinobacteria is a group of Gram-positive, spore-forming, chiefly filamentous bacteria with high GC content, that are widely distributed in soil. They can produce abundant bioactive secondary metabolites which contribute to about half of clinically-used antibiotics and important pharmaceutical agents (1–4). The biosynthesis of these metabolites depends on the available carbon, nitrogen and phosphate in bacteria, especially the nitrogen, which acts as an essential element, plays a pivotal role in bacterial survival and growth (5–7). To acquire more available nitrogen source from the constantly changing environment, bacteria have developed various and complex transcription regulatory systems to sense internal and external nitrogen status, by coordinating genes involved in nitrogen metabolism (5,8).

Two component systems (TCSs) have been identified as one of the simplest signal transduction systems. One TCS typically includes a membrane-associated histidine kinase (HK) and a cognate intracellular response regulator (RR) (9–11). RRs are involved in regulation of many behaviors, including metabolism, motility, quorum sensing, virulence, and antibiotic resistance(12–15). The OmpR/PhoB subfamily proteins are the largest group of RRs identified in bacteria. In low GC content enteric bacteria, the two component system NtrB-NtrC is adopted to regulate gene expression of nitrogen metabolism and have been well investigated (16,17). While in high GC content actinobacteria, metabolisms of nitrogen, carbon and phosphate are globally controlled by an OmpR/PhoB subfamily protein, GlnR, which is regarded as an orphan RR independent of Asp phosphorylation since none of its cognate HK has been identified so far (5,18). However, it was proposed to be possibly regulated by the post-translation modifications of Ser/Thr/Tyr phosphorylation or acetylation (19). GlnR and its homologs, comprising of an N-terminal receiver domain (GlnR-REC) and a C-terminal DNA binding domain (GlnR_DBD), are widely distributed in actinobacteria, such as the epidemic pathogen *Mycobacterium tuberculosis* (*M. tuberculosis*), the model organism *Streptomyces coelicolor* (*S. coelicolor*), the erythromycin-producing acetinobacteria *Saccharopolyspora erythraea* (*S. erythraea*), the rifamycin-producing industrial acetinobacteria *Amycolatopsis mediterranei* (*A. mediterranei*), and the soil dwelling saprophyte *Mycobacterium smegamatis* (*M. smegamatis*) (2,5,7,8,18,20).

Besides, GlnR has been characterized as a pleiotropic transcription regulator for expression of ~1000 genes to date (5,8,14,18). It can act as an activator or a repressor, mainly depending on its binding position in promoter DNA. The crystal structures of GlnR-RECs from *M. tuberculosis* and *A. mediterranei* have been solved and the relevant biochemical analyses demonstrated that GlnR can form functional dimer through its α4-β5-α5 interface from the GlnR-REC (20). Though the canonical GlnR binding *cis* element was firstly defined by Tiffert *et al* a decade ago (21), many trials were unsuccessful on acquiring the crystal structures of full-length GlnR, or GlnR_DBD, or GlnR_DBD in complex with their cognate *cis*-element DNA from different kinds of actinobacteria. It was probably due to the flexible loop connecting the N-terminal domain and the C-terminal domain of GlnR, which causes improper crystal packings.

The conserved GlnR *cis* element comprises two 22-bp GlnR boxes separated by six nucleotides. One GlnR box is composed of one “a1 site” of “gTnAc” and one “b1 site” of “GaAAC”, the other GlnR box includes one “a2 site” of “gTnAc” and one b2 site of “GaAAC” (21,22). However, in recent years, more and more GlnR target genes and their corresponding *cis*-elements have been identified, especially some atypical GlnR binding *cis*-elements. The *amtB* promoter from *S. coelicolor* was characterized to have three GlnR binding boxes consisting of a3-b3, a1-b1 and a2-b2 sites. The *nas* operon promoter of *A. mediterranei*, was identified to contain the previously deduced 22-bp GlnR binding consensus sequences a1-b1, a2-b2 sites (22). However, *in vitro* biochemical and *in vivo* mutational assays showed that only three of the above four GlnR binding sites (a1-b1 and a2-b2 sites) are essential for GlnR-dependent transcription activation (23), suggesting that the GlnR binding *cis*-element and GlnR-dependent transcription activation are complicated and elusive, and more experiments need to be performed to get a better understanding of the molecular mechanism.

Transcription initiation is a main step for gene expression and a predominant target for transcription regulation in bacteria. It usually includes assembly of the multi-subunit RNA polymerase (α_2_ββ’ωσ, RNAP) holoenzyme and the promoter DNA, isomerization of RNA polymerase-promoter closed complex (RPc), and subsequently formation of a catalytically competent RNA polymerase-promoter open complex (RPo) (24–26). As to promoters containing non-optimal consensus elements (−35 element and −10 element), various transcription activators will cooperate together with RNAP to form transcription activator-dependent transcription activation complexes (TACs) (27). So far, several classic bacterial transcription activators have been well studied, such as CAP and SoxS, which has only one cognate binding box located at different sites of promoter DNA (27–34). However, the underlying transcription activation mechanisms on promoters harboring two or three binding boxes as GlnR still remain unexplored.

In this report, we determined a crystal structure of GlnR_DBD in complex with its regulatory *cis*-element DNA (dsDNA), and a cryo-EM structure of GlnR-dependent transcription activation complexes (GlnR-TAC) comprising of *M. tuberculosis* RNA polymerase (RNAP), *M. tuberculosis* GlnR (GlnR) protein, and a promoter containing four well-characterized conserved GlnR binding sites. These structures defined the precise interactions between GlnR and its conserved *cis*-element DNA, revealed how GlnR tetramer initiates transcription through specific interactions with promoter DNA and σ^A^R4, αNTD, β flap from RNAP holoenzyme. Particularly, four GlnR_DBDs synergistically bind around the upstream and downstream of −35 element in a head-to-tail manner, two GlnR-RECs efficiently connect αNTD with GlnR_DBD located at a1 site, further stabilizing the GlnR-TAC complex. Additionally, one conserved C-terminal domain of RNAP alpha subunit (αCTD) cooperates with GlnR further stabilize the complex. In summary, our results here offer a distinctive molecular mechanism for actinobacteria-derived GlnR-dependent transcription regulation, and reveal a globally novel mode of bacterial transcription regulation.

## Materials and Methods

### Plasmids and DNA preparation

To construct the pET28a-*glnR* plasmid, *glnR* encoding an N-terminal His6 tagged *M. tuberculosis* GlnR under the control of T7 promoter was synthesized and recombined into pET28a by Sangon Biotech, Inc. Plasmids harboring *GlnR* amino acid substitutions (pET28a-*glnR derivatives*) were constructed using site-directed mutagenesis (QuikChange Site-Directed Mutagenesis Kit, Agilent, Inc.).DNA fragments consisting of different canonical GlnR-dependent promoters (*narG* whose downstream genes encoding *M. tuberculosis* nitrate reductases; *amtB6* and *amtB4* whose downstream genes encoding the ammonium transporter), followed by an RNA aptamer (Mango III) coding sequence were amplified by *de novo* PCR (22,35–38), and purified using the QIAquick PCR Purification Kit (Qiagen, Inc.), respectively. Promoter sequences and conserved elements are shown in **Supplementary Figure S1**. *amtB6* DNA, *amtB4* DNA*, amtB2* DNA are *amtB* mango DNA containing 6 GlnR binding sites (a3-b3, a1-b1 and a2-b2), 4 GlnR binding sites (a1-b1 and a2-b2), and 2 GlnR binding sites (a2-b2), respectively (**Supplementary Figure S1**). Primers used in this study are shown in **Supplementary Table S1**.

### Purification of *M. tuberculosis* GlnR and *S. erythraea* GlnR-DBD

Plasmids of pET28a-his TEV-*glnR or* pET28a-his TEV-*glnR* derivative was firstly transformed into expression strain BL21(DE3) (Invitrogen, Inc.). then single colonies of the positive transformants were inoculated and amplified with 5 L LB broth supplemented with 50 μg/ml kanamycin at 37 °C with shaking. Once OD_600_ of the cultures reached about ~0.8-1.0. GlnR expression was induced by adding 0.5 mM IPTG, and cultures were incubated for another 15 h at 20 °C. After being harvested by centrifugation (5,500 g; 15 min at 4 °C), the cell pellets were resuspended in 20 mL buffer A (20 mM Tris-HCl, pH 8.0, 0.2 M NaCl, 5% glycerol), and lysed using an ATS AH-10013 cell disrupter (ATS, Inc.). After centrifugation at 13,000 g for 30 min at 4 °C, the supernatant was loaded onto a 5 mL column of Ni-NTA agarose (Qiagen, Inc.) pre-equilibrated with buffer A. The column was washed with 25 mL buffer A containing 25 mM imidazole and eluted with 30 mL buffer A containing 200 mM imidazole. Subsequently the elutes were concentrated and applied to a 120 mL HiLoad 16/600 Superdex 75 column (GE Healthcare, Inc.) equilibrated with buffer B (20 mM Tris-HCl, pH 8.0, 75 mM NaCl, 5 mM MgCl_2_), and the column was eluted with the same buffer for a column volume. The targeted fractions containing GlnR identified by SDS-PAGE were pooled and stored at –80 °C. Yield was ~3.0 mg/L, and purity was > 95%. GlnR derivatives or *Saccharopolyspora erythraea* GlnR-DBD (*Sae*GlnR-DBD) were prepared as described above. In order to remove the N-terminal his tag, the elutes of GlnR147 (his-TEV GlnR C60SL147C) from Ni-NTA column were cleaved with recombinant tobacco-etch virus protease (Life Technologies) overnight at 4 °C, subsequently exchanged by buffer A, and applied onto a second Ni-NTA column to remove the residual his tagged GlnR and protease. The flow-through sample from the second Ni-NTA column were finally concentrated and purified by HiLoad 16/600 Superdex 75 column as GlnR.

### Purification of *M. tuberculosis* RNAP

*M. tuberculosis* RNAP was prepared from cultures of *E. coli* strain BL21(DE3) (Invitrogen, Inc.) co-transformed with plasmids of pACYC Duet-*rpoA-rpoD*, pCDF-*rpoZ* and pET Duet-*rpoB-rpoC*, and purified as described with some modifications (39). Single colonies of the resulting transformants were used to inoculate 100 mL LB broth containing 35 μg/mL chloramphenicol, 100 μg/mL ampicillin, 50 μg/mL streptomycin, and cultures were incubated for 16 h at 37 °C with shaking. Subsequently, cultures were amplified by transferring every 10 mL aliquots into 1 L LB broth containing the same antibiotics, and incubated at 37 °C with shaking. When OD_600_ reached ~0.8-1.0, cultures were induced by addition of 0.5 mM IPTG, and incubated for 15 h at 20 °C. Then cells were harvested by centrifugation (5,000 g; 15 min at 4 °C), resuspended in 20 ml lysis buffer A, lysed using a JN-02C cell disrupter (JNBIO, Inc.) and centrifugated at 13,000 g for 30 min under 4 °C condition. Then the supernatants were precipitated by poly(ethyleneimine) to a ratio of 0.7% (m/v), washed three times by buffer C (10 mM Tris-HCl, pH 8.0, 0.5 M NaCl, 1 mM EDTA, 5% glycerol), extracted by buffer D (10 mM Tris-HCl, pH 8.0, 1.0 M NaCl, 1 mM EDTA, 5% glycerol), and subsequently by precipitated by ammonium sulfate to a ratio of 30.0% (m/v). The pellets were resuspended with buffer A and loaded onto a 10 mL column of Ni-NTA agarose (Qiagen, Inc.) equilibrated with buffer A. The column was washed with 50 mL buffer A containing 20 mM imidazole and eluted with 50 mL buffer A containing 0.3 M imidazole. The eluate was diluted with buffer E (20 mM Tris-HCl, pH 7.9, 5 % glycerol, 1 mM EDTA and 1 mM DTT) and loaded onto a Mono Q 10/100 GL column (GE Healthcare, Inc.) equilibrated in buffer E and eluted with a 160 mL linear gradient of 0.3–0.5 M NaCl in buffer E. Fractions containing *M. tuberculosis* RNAP were pooled and applied to a 120 mL HiLoad 16/600 Superdex 200 column (GE Healthcare, Inc.) equilibrated in buffer B, and the column was eluted with the same buffer. Fractions containing *M. tuberculosis* RNAP were pooled and stored at −80°C. Yield was ~0.5 mg/L, and purity was > 95%.

### Crystallization of *Sae*GlnR-DBD complexed with its *cis*-element DNA

To assemble the binary complex of *Sae*GlnR-DBD bound to its *cis*-element DNA, two strands of a designated conserved *cis*-element DNA (21 bp long, 5’-AC**GTAAC**ATCGCG**GTAAC** A-3’) consisting of two copies of GlnR binding sites (a site and b site shown in bold) (**Figure 1A**) were synthesized and gel purified by Sangon Biotech, Inc. Firstly, template strand oligonucleotide and nontemplate strand oligonucleotide of this blunted DNA were dissolved into nuclease-free water to 1 mM, and annealed at the ratio of 1:1 in the annealing buffer (10 mM Tris-HCl, pH 7.9, 0.2 M NaCl). *Sae*GlnR_DBD and DNA were incubated in a molar ratio of 2: 1.1 at 4 °C for 1 h. Then the sample were centrifugated and applied to robotic crystallization trials by using a Gryphon liquid handling system (Art Robbins Instruments), commercial screening solutions (Emerald Biosystems, Hampton Research, and Qiagen), and the sitting-drop vapor-diffusion technique (drop: 0.2 μL sample mixed with 0.2 μL screening solution; reservoir: 60 μL screening solution; 22 °C). A total of 1000 conditions were screened and brick-shaped crystals appeared within 1 week. Through optimization by hanging-drop vapor-diffusion technique at 16 °C, high-quality crystals were obtained under the condition of 0.1 M Bis-tris pH 5.7, 25% PEG3350 in 1 week. Crystals were transferred to reservoir solution containing 18% (*v/v*) (*2R,3R*)-(-)-2,3-butanediol (Sigma-Aldrich) and flash-cooled with liquid nitrogen.

**Figure 1.**
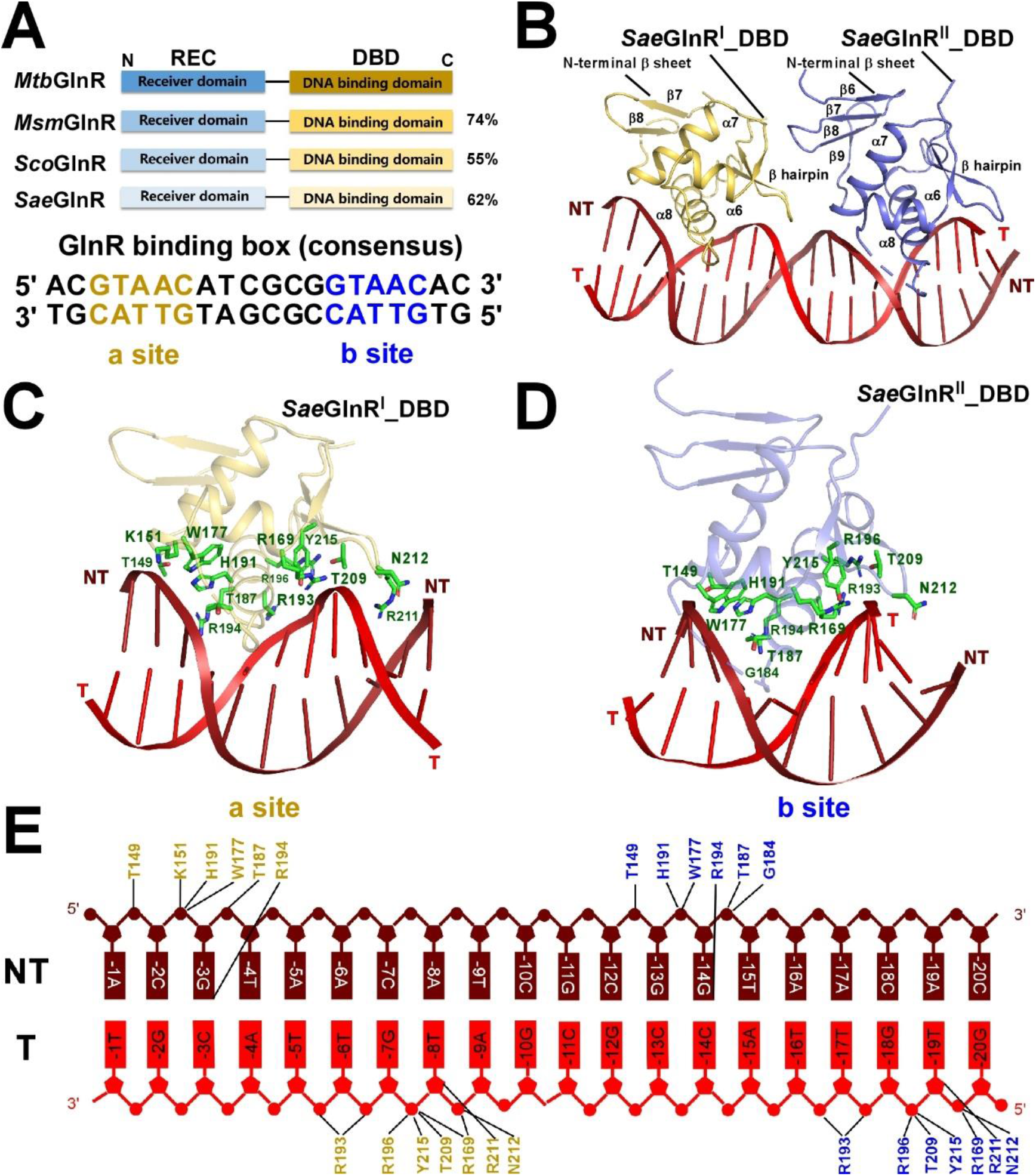
The co-crystal structure of *Sae*GlnR bound to the designed conserved consensus promoter DNA. **(A)** Domain architectures of *M. tuberculosis* GlnR (*Mtb*GlnR), *M. smegmatis* GlnR (*Msm*GlnR), *S. coelicolor* GlnR (*Sco*GlnR), and *S. erythraea* GlnR (*Sae*GlnR) (top panel); the sequences of designed conserved consensus promoter DNA for *Sae*GlnR protein with the a- and b-site sequences highlighted in orange and blue, respectively (bottom panel). Domains of REC (N-terminal receiver domain) and DBD (C terminal DNA binding domain) are individually labeled above the architectures. **(B)** Co-crystal structure of *Sae*GlnR^I^_DBD, *Sae*GlnR^II^_DBD in complex with the designed conserved consensus promoter DNA. *Sae*GlnR^I^_DBD and *Sae*GlnR^II^_DBD are represented as orange and blue cartoon, respectively. NT, non-template-strand promoter DNA (in dark red cartoon); T, template-strand promoter DNA (in red cartoon. **(C-D)** Detailed interactions between *Sae*GlnR and the designed conserved consensus promoter DNA. Residues from *Sae*GlnR involved in interacting with a-site or b-site of promoter DNA are shown in green sticks. **(E)** Summary of the interactions between the promoter DNA and *Sae*GlnR. Black lines indicate residues that contact DNA.

### Crystal data collection and structure determination

By using the cryo-cooled crystals, diffraction data of *Sae*GlnR-DBD complexed with DNA were collected at Shanghai Synchrotron Radiation Facility (SSRF) beamline 17U, followed by data processing with HKL2000 (40). Finally, the structure was solved by molecular replacement with Molrep (41) using the structure of PhoB effector domain in complex with pho box DNA (PDB 1GXP) (42) as the search model. The model of *Sae*GlnR-DBD complexed with DNA was built in Coot (43) and refined in Phenix (44). Structure data collection and refinement statistics are listed in **Table 1.**

**Table 1.**
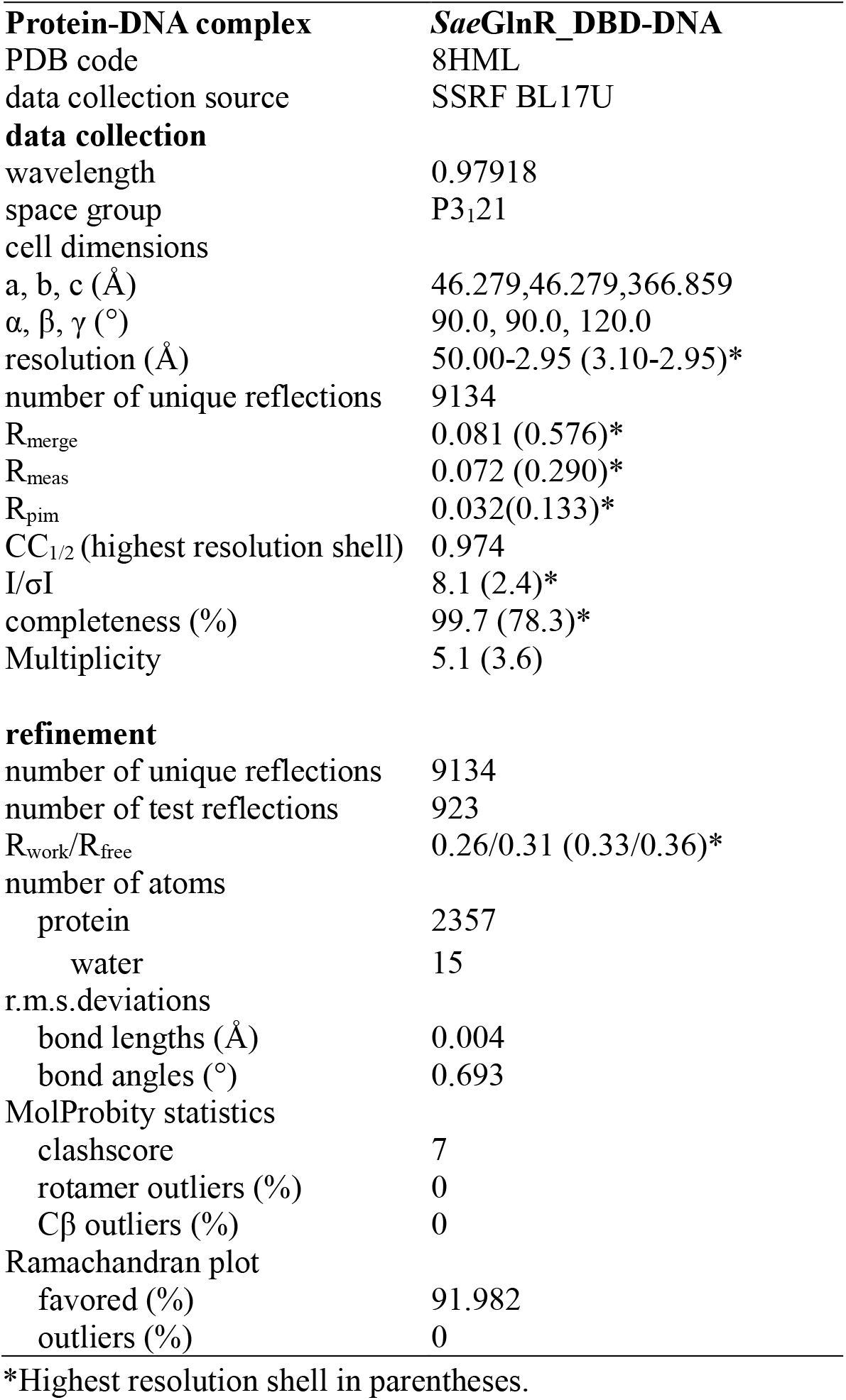
Structure data collection and refinement statistics.

### Assembly of *M. tuberculosis* GlnR-TAC

DNA oligonucleotides used to assemble *M. tuberculosis* GlnR-TAC were synthesized and gel purified by Sangon Biotech, Inc. Template strand DNA (*amtB* scaffold_T) and nontemplate strand DNA (*amtB* scaffold_NT) of this *amtB* scaffold (**Figure 2A**) were firstly dissolved into nuclease-free water to 1 mM, and annealed at the ratio of 1:1 in the annealing buffer (10 mM Tris-HCl, pH 7.9, 0.2 M NaCl). Then,*M. tuberculosis* GlnR-TAC was assembled by incubating *M. tuberculosis* RNAP, *amtB* scaffold, and *M. tuberculosis* GlnR in a molar ratio of 1: 1: 10 at 4 °C overnight. The sample was then applied onto a 120 mL HiLoad 16/600 Superdex 200 column (GE Healthcare, Inc.) equilibrated in buffer B, and the column was eluted with the same buffer. The elutes were identified by SDS-PAGE and electrophoretic mobility shift assay (EMSA). Finally, fractions containing right assembled *M. tuberculosis* GlnR-TAC were concentrated to 40 mg/mL using Amicon Ultra centrifugal filters (10 kDa MWCO, Merck Millipore, Inc.). Sequences of the above *amtB* oligonucleotides are shown in **Supplementary Table S1**.

**Figure 2.**
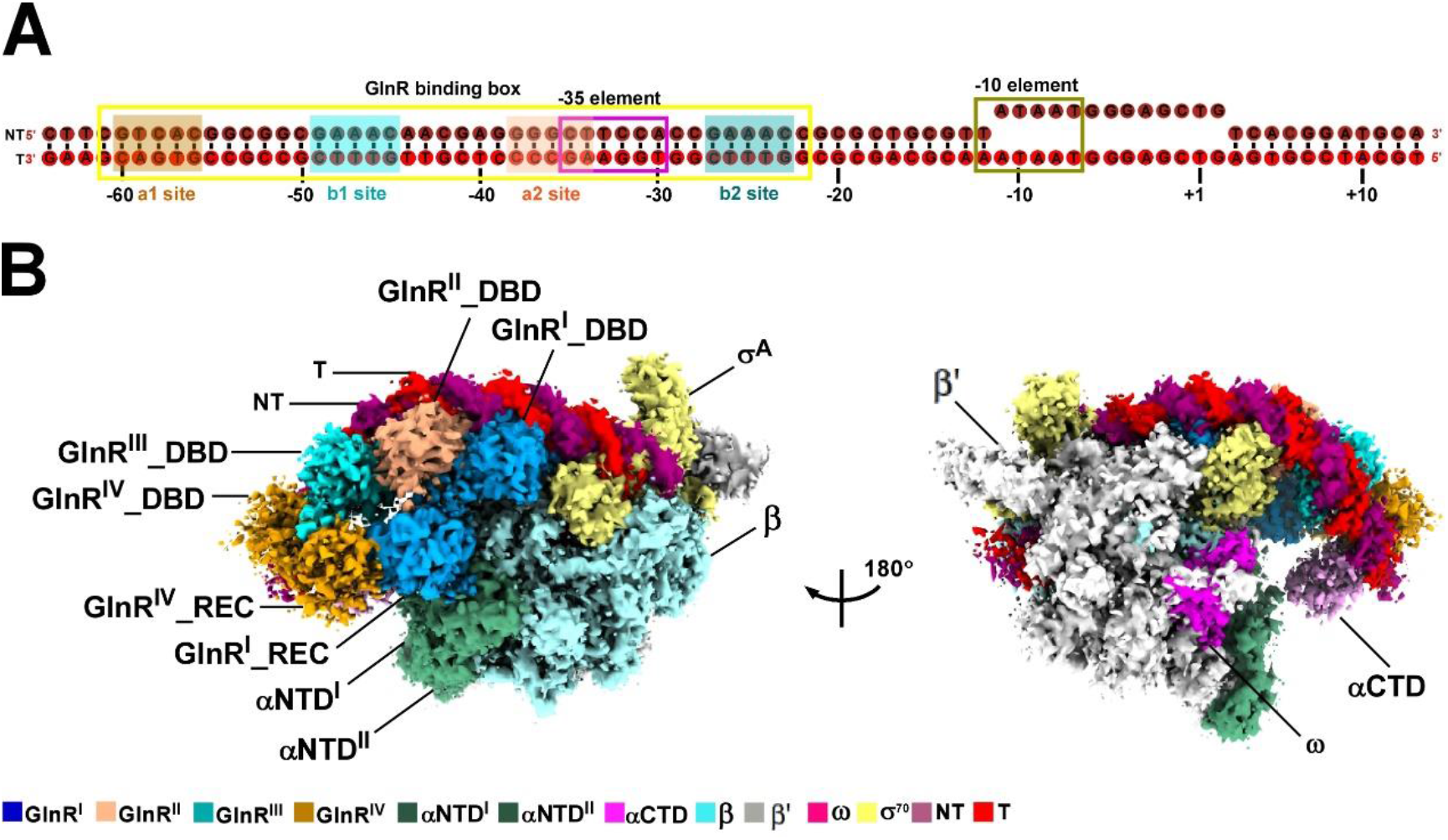
The overall structure of *M. tuberculosis* GlnR-TAC. **(A)** DNA scaffold used in structure determination of *M. tuberculosis* GlnR-TAC (top panel). NT, non-template-strand promoter DNA; T, template-strand promoter DNA. GlnR binding box is framed in yellow color, with a1 site, b1 site, a2 site, and b2 site shaded in orange, cyan, light orange and blue colors, respectively. The −35 element and −10 element are enclosed with pink and brown frames, respectively. **(B)** Two views of the cryo-EM density map of *M. tuberculosis* GlnR-TAC (bottom panel). The EM density maps and cartoon representations of *M. tuberculosis* GlnR-TAC are colored as indicated in the color key.

### Cryo-EM grid preparation

C-flat grids (CF-1.2/1.3 400 mesh holey carbon grids; Protochips, Inc.) were glow-discharged for 60 s at 15 mA. Once being incubated with CHAPSO to a final concentration of 8 mM (Hampton Research Inc.), 3 μL of the purified *M. tuberculosis* GlnR-TAC was applied onto the grids, blotted with Vitrobot Mark IV (FEI), and immediately plunge-frozen in liquid ethane with 95 % chamber humidity at 10 °C.

### Cryo-EM data collection and processing

Cryo-EM data of *M. tuberculosis* GlnR-TAC were collected by using a 300 kV Titan Krios (FEI, Inc.) equipped with a K3 Summit direct electron detector. A total of 1,200 images were recorded with EPU in counting mode with a pixel size of 1.2 Å, a dose rate of 10 e/pixel/s, and an electron exposure dose of 50 e/Å^2^. Movies were recorded for 8.38 s and defocus range varied between −1.4 μm and −2.2 μm. Subframes of individual movies were aligned using MotionCor2 (45), and contrast-transfer-function for each summed image was estimated using CTFFIND4. From the summed images, approximately 10,000 particles were manually picked and subjected to 2D classification in RELION 3.1 (46). The corresponding distinct 2D classes were used as templates for particle auto-picking. 202,298 particles were auto-picked, manually inspected, and subjected to further 2D classification by specifying 100 classes. By removing the poorly populated classes, 118,770 particles were subjected to 3D classification in RELION by using a map of *M. tuberculosis* RPo (PDB ID: 6VVY) (47) low-pass filtered to 40 Å resolution as a reference. Then particles in Class 2 of the first three resulted 3D classes were 3D-classified, following by class 1 running into 3D auto-refinement. The selected 54,432 particles were further re-extracted, CTF-refined, Bayesian polished, 3D auto-refined and post-processed in RELION (**Supplementary Figure S5**). The Gold-standard Fourier-shell-correlation analysis indicated a mean map resolution of 3.75 Å of *M. tuberculosis* GlnR-TAC (**Supplementary Figure S6**).

### Model building and refinement

The model of *M. tuberculosis* RPo (PDB ID: 6VVY) (47) and the co-crystal structure of *Sae*GlnR_DBD bound to its *cis*-element DNA were manually fitted into the cryo-EM density map of GlnR-TAC by Chimera (48). Data was further calculated and validated by rigid body and real-space refinement in Coot and Phenix. Structures were analyzed with PyMOL (49) and Chimera.

### *In vitro* transcription assay

*In vitro* transcription assays were performed in transcription buffer (40 mM Tris-HCl, pH 8.0, 50 mM NaCl, 10 mM MgCl_2_, 5% glycerol) by using 96-well microplates (Corning incorporated, USA). Reaction mixtures (80 μL) contained: 0.1 μM *M. tuberculosis* RNAP, 30 nM mango-ended DNA, 0 or 8 or 16 μM *M. tuberculosis* GlnR or its derivatives, 0.1 mM NTP mix (ATP, UTP, GTP, and CTP), and 1 μM TO1-Biotin. Once *M. tuberculosis* RNAP and DNA were incubated for 10 min at 37 °C, *M. tuberculosis* GlnR was added into the mixture and incubated for 20 min at 37 °C. Subsequently, NTP mix and TO1-biotin were added and the mixture was incubated for another 30 min at 37 °C. Finally, fluorescence emission intensities were measured using a multimode plate reader (EnVision, PerkinElmer Inc.; excitation wavelength = 510 nm; emission wavelength = 535 nm). Relative transcription activities of GlnR derivatives were calculated using:

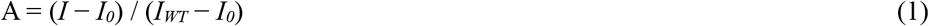

where *I_WT_* and *I* are the fluorescence intensities in the presence of GlnR and GlnR derivatives; *I_0_* is the fluorescence intensity in the absence of GlnR.

### Electrophoretic mobility shift assay

Electrophoretic mobility shift assay (EMSA) was carried out in EMSA buffer (40 mM Tris-HCl, pH 8.0, 100 mM NaCl, 10 mM MgCl_2_, 5 % glycerol). Components and final concentration of the reaction mixture (20 μL) are as following: 0 or 8 μM *M. tuberculosis* GlnR (or GlnR derivatives), 100 nM *M. tuberculosis* RNAP, 30 nM promoter *amtB4* DNA. RNAP was firstly incubated with DNA for 10 min at 37 °C, then incubated with GlnR or GlnR derivatives for 20 min at 37 °C. After addition of 0.03 mg/ml heparin and incubation for 2 min at 22 °C, the reaction mixtures were applied to 5% polyacrylamide slab gels (29:1 acrylamide/bisacrylamide), electrophoresed in 90 mM Tris-borate, pH 8.0, and 0.2 mM EDTA, and stained with 4S Red Plus Nucleic Acid Stain (Sangon Biotech, Inc.) according to the procedure of the manufacturer.

### Single-molecule fluorescence assay

To prepare a fluorescence detectable complex of *M. tuberculosis* GlnR-TAC, the purified *M. tuberculosis* GlnR147 was labeled with Alexa647 to generate GlnR-Alexa647, and DNA (amtB+2Cy3) was constructed to carry a Cy3 label at position +2 of the non-template DNA (amtBpNT+2Cy3). GlnR-TAC was assembled by incubating *M. tuberculosis* RNAP (110 nM) and amtB+2Cy3 DNA (56 nM) in a final volume of 10 μl for 30 min at 37 °C with shaking (550 rpm). Then, GlnR-Alexa647 was added to a final concentration of 1.8 μM in a final volume of 50 μl, and incubated for 30 min at 37 °C with shaking (550 rpm). Subsequently, the sample can be subjected to single-molecule fluorescence measurements. Details are described in the **Supplementary Materials and Methods**.

Single-molecule fluorescence assay was performed under a home-made TIRF-based microscope equipped with two OPIS lasers (532 nm and 640 nm, Coherent). To illuminate fluorescent dyes, an alternative laser excitation (ALEX) module was implemented to operate at 10 mW and 30 mW for these two lasers, respectively. Fluorescence was imaged by an objective (Apo TIRF 100x, 1.49NA, Olympus), split into two channels and then data collected on an EMCCD (iXon Ultra 897, Andor). A flow chamber was sandwiched with a glass surface with two drilled holes, a parafilm layer and an undrilled glass surface. Glass surfaces were prepared as previously described (33). Prior to use, the flow chamber was incubated with 10 μg/ml streptavidin (ThermoFisher Scientific) in 1x PBS for 10 min followed by flushing away free streptavidin. Biotin anti-His tag antibody (abcam) was then flushed in the channel at 10 μg/ml concentration in 1x PBS buffer for 10 min and then flushed away free antibodies. The complex of prepared *M. tuberculosis* GlnR-TAC was tethered to the glass surface through these anti-His tag antibodies. Fluorescence images were collected at an acquisition rate of 20 Hz in transcription buffer containing 0.5 mg/ml BSA, 1 mg/ml glucose oxidase, 0.4 mg/ml catalase, 0.8% glucose and 1 mM Trolox (Sigma-Aldrich). Raw fluorescence data *I_D_* and *I_A_* were extracted from the donor and acceptor channels respectively and background corrected. The number of steps that Alexa647 photobleached was analyzed manually.

### Qualification and statistical analysis

The FSC cut-off criterion of 0.143 (50) was used to calculate Fourier shell correlations (FSC) of GlnR-TAC (**Supplementary Figures S6B, S6E**). To assess the data of transcription assays (**Figures 3E, 4D and 5G**), the mean values and their corresponding standard errors from three independent measurements were analyzed and displayed. The local resolution of the cryo-EM maps (**Supplementary Figures S6C and S6D**) was estimated using blocres (51). PHENIX (44) was also used for quantification and statistical analyses during model refinement and validation of GlnR-TAC.

**Figure 3.**
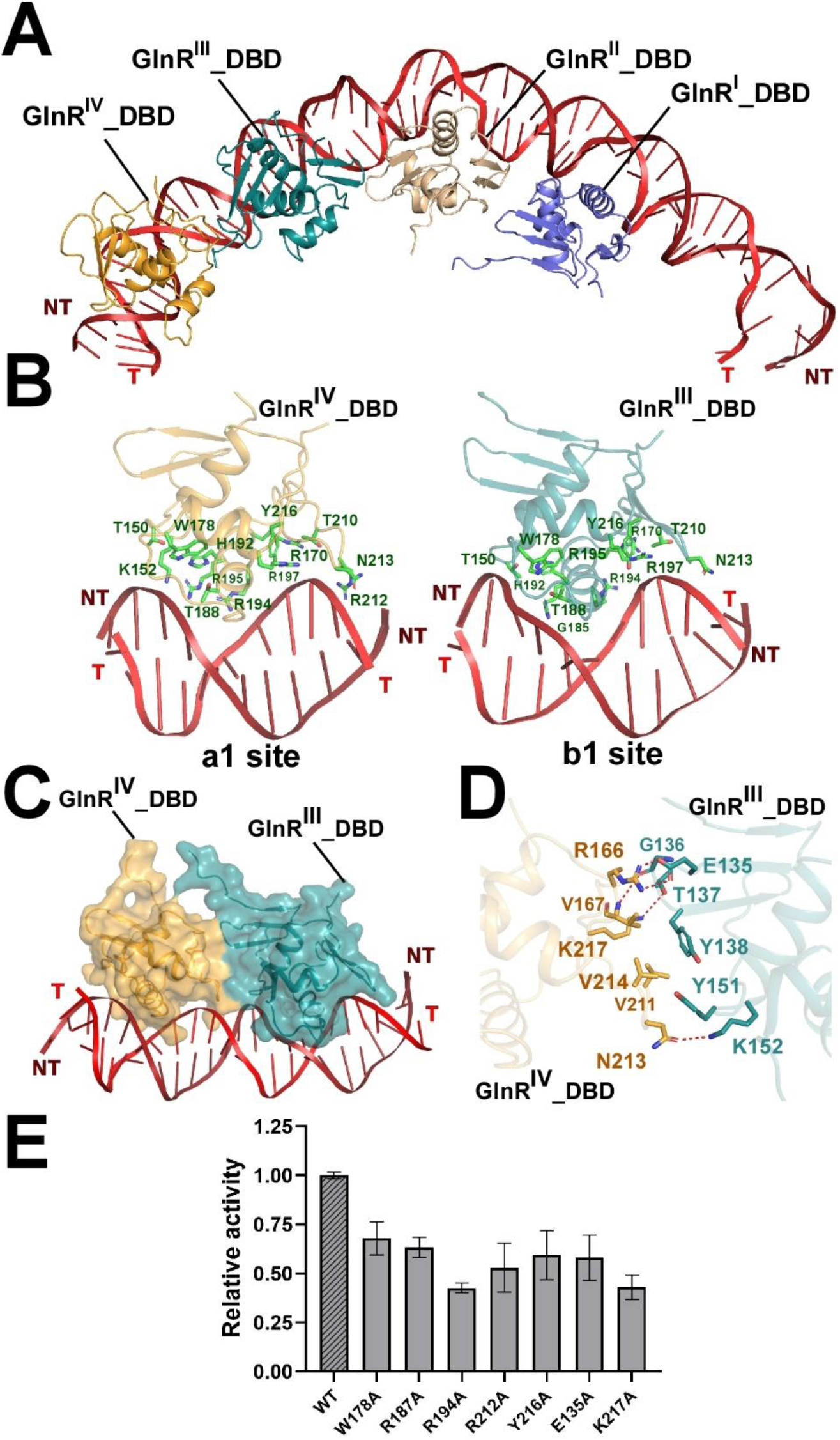
Four GlnR molecules engage promoter DNA in *M. tuberculosis* GlnR-TAC. **(A)** Relative locations of *M. tuberculosis* GlnR^I^_DBD, GlnR^II^_DBD, GlnR^III^_DBD, and GlnR^IV^_DBD located at the upstream double-stranded DNA. **(B)** Detailed interactions between *M. tuberculosis* GlnR^IV^_DBD, GlnR^III^_DBD and their corresponding GlnR binding sites. **(C-D)** The relative locations and detailed interactions between GlnR^IV^_DBD and GlnR^III^_DBD bound to the promoter DNA. Salt-bridges are shown as red dashed lines. **(E)** Substitutions of GlnR residues involved in promoter engagement reduced *in vitro* transcription activity. Colors are shown as in **Figure 2**.

**Figure 4.**
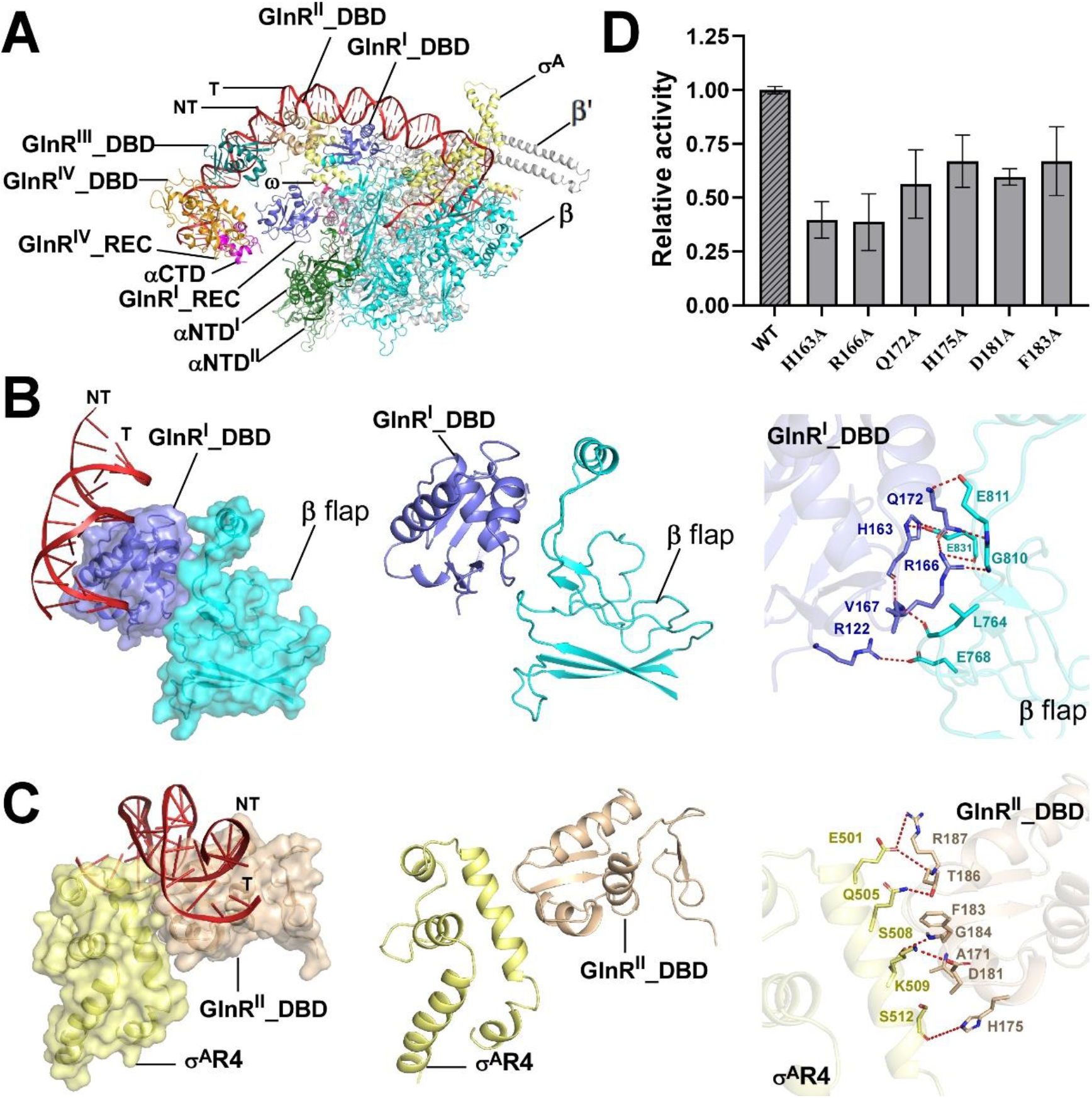
The critical protein-protein interactions between *M. tuberculosis* GlnR and domains of RNAP β flap, σ^A^R4. **(A)** The structure model of *M. tuberculosis* GlnR-TAC. **(B)** Relative locations of *M. tuberculosis* GlnR^I^_DBD, RNAP β flap, and the double-stranded DNA of GlnR binding site b2 (left, middle panel); Detailed interactions between *M. tuberculosis* GlnR^I^_DBD and RNAP β flap (right panel). **(C)** Relative locations of *M. tuberculosis* GlnR^II^_DBD, RNAP σ^A^R4, and the double-stranded DNA of GlnR binding site a2 (left, middle panel); Detailed interactions between *M. tuberculosis* RNAP σ^A^R4 and GlnR^II^_DBD. Polar interactions are shown as red dashed lines. **(D)** Mutational effects of the key GlnR residues involved in the GlnR-β flap, GlnR-σ^A^R4 interfaces by *in vitro* transcription assays. Colors are shown as in **Figure 2**.

**Figure 5.**
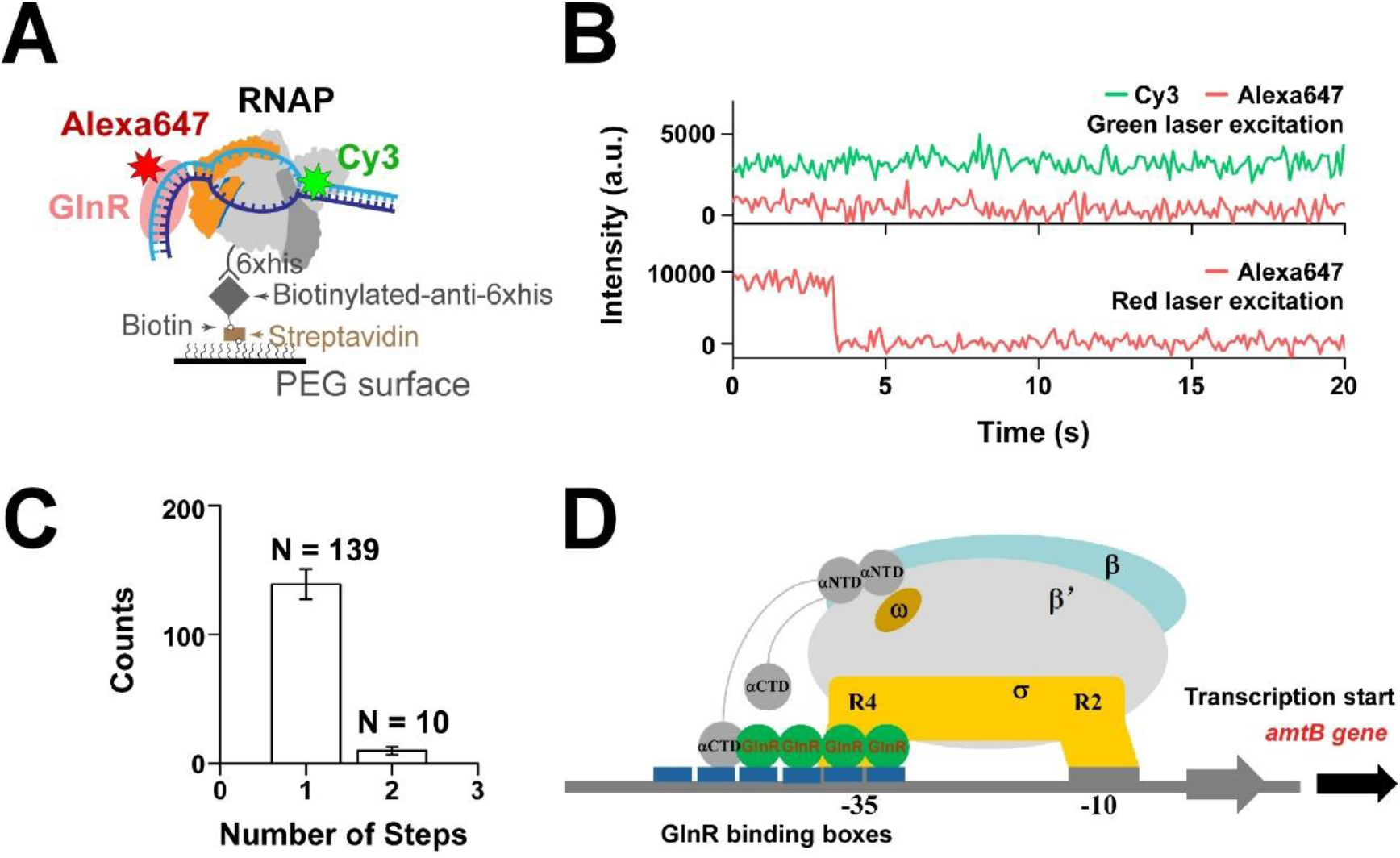
Four GlnR molecules kinetically activates formation of a competent GlnR-TAC. **(A)** Scheme of single-molecule fluorescence assay. The complex of GlnR-TAC was assembled with Cy3 labeled DNA (+2 position, amtB+2Cy3) and Alexa647 labeled GlnR147 (GlnR-Alexa647) and immobilized on a glass surface. Fluorescence of these two dyes were collected via an ALEX module and analyzed. **(B)** A typical trajectory displaying Cy3 and Alexa647 fluorescence under green laser excitation and Alexa647 fluorescence under red laser excitation. Under green laser excitation, Alexa647 fluorescence can hardly be observed suggesting no obvious FRET between Cy3 and Alexa647 in the complex. Under red laser excitation, one step of Alexa647 photobleaching was observed. **(C)** Statistics representing Alexa647 photobleaching events with one step (N = 139) and two steps (N = 10). The mean number of steps is calculated as 1.07 ± 0.02 (SEM). **(D)** Proposed working model for GlnR-dependent transcription activation. Four GlnR molecules preferentially and kinetically activates formation of a stable and competent GlnR-TAC. GlnR binding boxes are presented in dark blue.

## Results

### Co-crystal structure of *Sae*GlnR_DBD bound to its *cis*-element DNA

To understand the structural basis of GlnR’s DNA binding specificity to conserved GlnR binding *cis* element, we tried to crystallize full-length GlnRs or GlnR-DBDs from several representative actinobacteria sharing high similarity (**Figure 1A; Supplementary Figure S2**). Fortunately, we finally succeeded in determining a 2.95 Å-resolution crystal structure of *S. erythraea* GlnR_DBD (*Sae*GlnR_DBD) bound to a designated conserved *cis*-element DNA which includes two copies of GlnR binding sites (a site and b site with a consensus sequence of “GTAAC”) (**Figure 1A**). The statistics of data collection and model refinement are summarized in **Table 1**. The GlnR_DBD, consisting of residues 122-221 which includes three helices and two β-sheets consisting of seven β strands, forms a typical winged-helix-turn-helix structural topology of the OmpR/PhoB subfamily proteins (42,52–55) (**Figure 1B**). In the co-crystal structure, there are two GlnR_DBD molecules and one blunt-ended dsDNA molecule in one asymmetric unit, with two GlnR_DBD molecules binding in tandem to the dsDNA. The 2Fo-Fc electron density map shows unambiguous, sharp densities for all nucleotides and residues from DNA and proteins, respectively. The α8 helices from two GlnR_DBD molecules contact the major grooves of dsDNA while the C-terminal β-hairpins interact with the adjacent minor grooves in a head-to-tail orientation, respectively. This head-to-tail domain arrangement for DNA recognition is similar to those for other OmpR/PhoB subfamily members like PhoB or PmrA (52,53). The specific DNA-protein recognition mainly includes direct hydrogen bond interactions, hydrophobic interactions, and Van der Waals forces (**Figure 1C-E**). As to a site from the dsDNA, *Sae*GlnR^I^_DBD residues T149, K151, H191, T187, and residues R169, R193, R196, T209, R211, N212, Y215 form direct hydrogen-bonds with the backbone phosphates of the −2C, −3G, −4T nucleotides from non-template and the 6T, −7G, −8T, −9A nucleotides from the template strand DNA, respectively (**Figure 1C, 1E**). By analogy, residues T149, H191, T187, G184, R194, R193, R196, T209, R169, N212, Y215 from *Sae*GlnR^II^_DBD also form direct hydrogen-bonds with the backbone phosphates of the −13G, −14G, −15T, −17T, −18G, −19T, −20G nucleotides from b site, respectively (**Figure 1D, 1E**). Each residue W177 from the two GlnR_DBD molecules interact with their corresponding −3G and −14G through hydrogen-bond interactions and Van der Waals forces. Residue R194 from each GlnR_DBD makes direct hydrogen-bond interactions with the corresponding −3G and −14G (**Figure 1C-E**).

### Overall structure of GlnR-TAC

To better understand the mechanism of GlnR-dependent transcription activation, we tried to obtain the GlnR-TAC structure, and then constructed one DNA scaffold harboring GlnR-dependent promoter *amtB* (from −64 to +13, positions numbered relative to the transcription start site) (**Figure 2A**) (22,23). The promoter includes two GlnR binding boxes with typical a1, b1, a2, b2 sites, a non-optimal −35 element, a consensus −10 element, a 13-nucleotide (nt) transcription bubble, and an 11-bp downstream DNA. SDS-PAGE analysis of the purified complex showed that *M. tuberculosis* RNAP holoenzyme and *M. tuberculosis* GlnR were included as expected, indicating of one well-assembled GlnR-TAC (**Supplementary Figure S3A, 3B**). Meanwhile, we constructed three fragments of promoter DNA containing different GlnR-dependent promoters followed by one *Mango III* sequence encoding a fluorogenic aptamer (**Supplementary Figure S1**). Upon addition of the purified *M. tuberculosis* GlnR, *M. tuberculosis* RNAP showed obvious transcription activities on previously identified *M. tuberculosis* promoter *narG* DNA, and also on the *amtB* DNA containing four GlnR-binding sites (*amtB4* DNA containing a1, b1, a2, b2 sites) or six GlnR-binding sites (*amtB6* DNA containing a1, b1, a2, b2, a3, b3 sites). As suggests that the proteins prepared were physiologically relevant and *M. tuberculosis* GlnR-TAC is functionally assembled (**Supplementary Figure S3C**).

We then collected cryo-EM datasets and determined one intact GlnR-TAC structure at a nominal resolution of 3.75 Å (**Figure 2B; Supplementary Figures S4 and S5; Table 2**). The electron densities for RNAP holoenzyme and the downstream DNA are unambiguous and each component from the structure of *M. tuberculosis* RPo (PDB ID: 6VVY) could be well fitted into the cryo-EM map (**Supplementary Figures S6**). The calculated local resolution is ~ 3.0–4.5 Å for the core RNAP, and ~5.5–7.5 Å for the peripheral αCTD and GlnR, indicating of their flexibility (**Supplementary Figures S5, S6C-F**). Consistent with the previous biochemical and genetic experiments (21–23), four GlnR protomers simultaneously associate with the two GlnR binding boxes of the promoter DNA in a head-to-tail fashion in the GlnR-TAC structure (**Figure 2–5; Supplementary Figures S7**). Each GlnR_DBD specifically contacts the a site and b site with its α8 helix and β–hairpin as those in the binary structure of *Sae*GlnR_DBD in complex with its *cis*-element DNA (**Figure 3A; Supplementary Figure S7**). Strikingly, different from the previously reported classic transcription activation complexes (27–29,31,33,34), GlnR^I^-DBD binds to the b2 site located downstream of the −35 element, and simultaneously interacts with β flap and σ^A^R4 of RNAP (**Figure 4B, 4C; Supplementary Figure S9)**. Additionally, the electron density of αCTD is unambiguous, showing αCTD engages around a1 site (**Supplementary Figure S10)**. These observations suggest that the structural and functional architecture of GlnR-TAC is in a novel transcription activation mode, which is distinct from the reported class I and II transcription activation complexes.

**Table 2.**
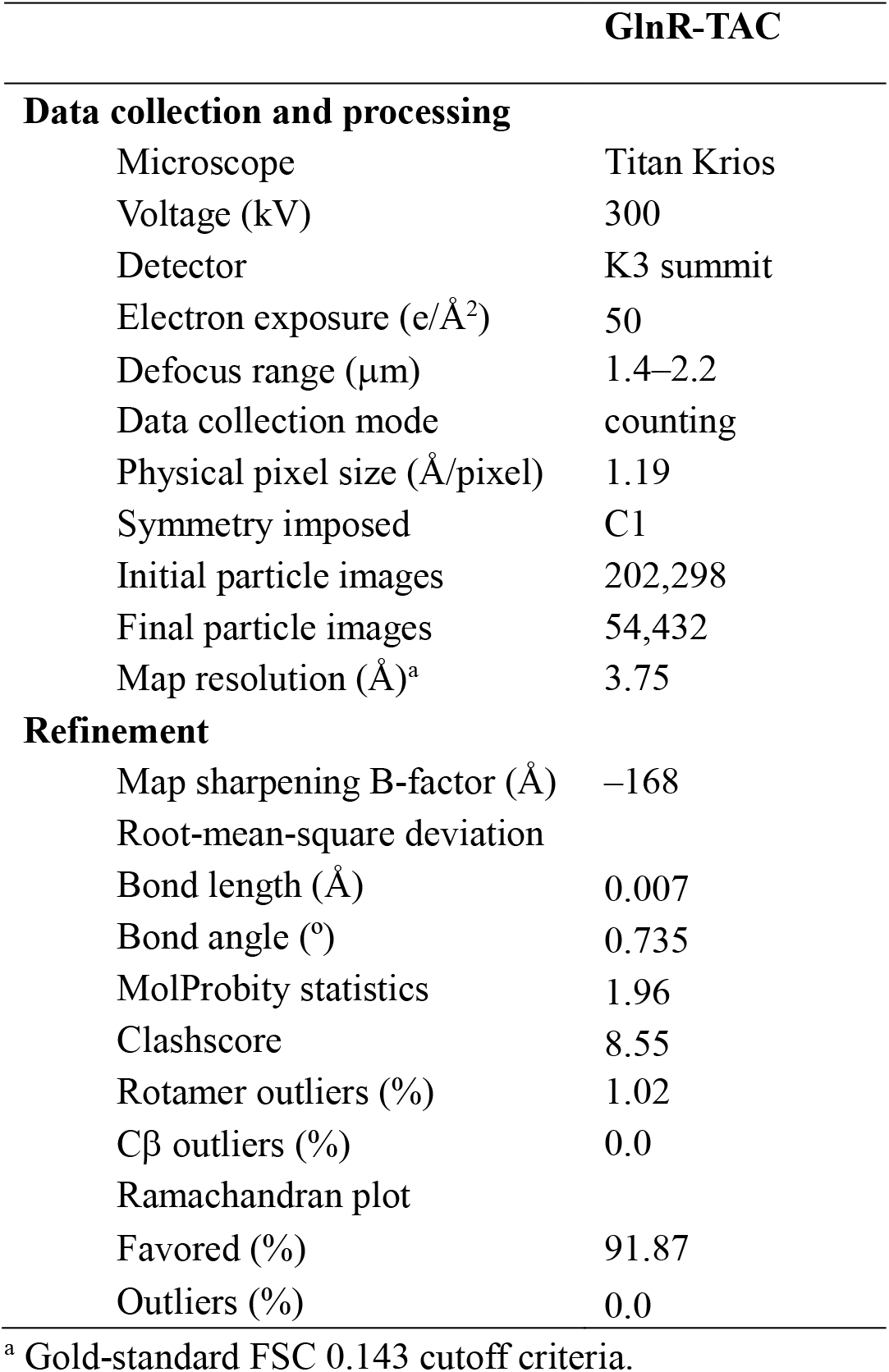
Single particle cryo-EM data collection, processing, and model building for *M. tuberculosis* GlnR-TAC.

### Four GlnR protomers coordinately engage promoter DNA in GlnR-TAC

Although previous studies suggested that the number of GlnR binding boxes might vary from one to three in the promoters of different target genes, the classic two GlnR binding boxes are found in the promoters of most GlnR target genes (21–23,38). To uncover the molecular mechanism, the *amtB* promoter, containing two typical GlnR boxes (including a1, b1, a2, b2 sites) was chosen to assemble GlnR-TAC. In good agreement with the previous reports, four *M. tuberculosis* GlnR protomers engage their corresponding GlnR binding sites located at the DNA major grooves using the α8 recognition helix from each GlnR_DBD in GlnR-TAC (**Figure 3A, 3B; Supplementary Figure S7A**). Binding of four GlnR protomers bend the linear dsDNA gently with a curvature of ~50° (**Figure 3A**), larger than that was observed in the PhoB_DBD-DNA complex containing two PhoB molecules (52). This may render GlnR easier to contact the core enzyme of RNAP, and further stabilize the transcription initiation complex. This binding mode of these *M. tuberculosis* GlnR_DBD to DNA shares great similarity with that in the co-crystal structure of *Sae*GlnR_DBD in complex with its conserved *cis*-element DNA. A cluster of residues consisting of T188, R194, R195, H192, R170, R197 from the α8 helix, T150, K152 and W178 from the linker connecting the α6 and α7 helices, Y216, T210, R212, N213 from the C-terminal β hairpin of GlnR^IV^_DBD make extensive contacts with the major and minor grooves of a1 site by salt bridges, hydrogen bonds, and van der Waals forces (**Figure 3B, left panel**). Analogously, residues T188, G185, R194, R195, H192, R170, R197, T150, K152, Y216, T210, R212, N213 from GlnR^III^_DBD also contribute to stabilization of the interface between GlnR^III^_DBD and b1 site DNA (**Figure 3B, right panel**). For GlnR^I^_DBD and GlnR^II^_DBD, similar interactions are included in the interfaces between GlnR_DBD and a2/b2 sites (**Supplementary Figure S7A**). Besides, interface between two adjacent GlnR_DBDs is stabilized by hydrogen bonds, hydrophobic interactions and π–π stacking forces, which is the same case between two *Sae*GlnR_DBDs (**Figure 3C, 3D; Supplementary Figure S7B-E)**. Consistently, substitutions of these residues implicated in the above interfaces resulted in defects in GlnR-dependent transcription activity and formation of GlnR-TAC as identified by our *in vitro* transcription assays and EMSA experiments (**Figure 3E; Supplementary Figure S9A and S9B**), reflecting their physiological importance in retaining such a special “head-to-tail” mode of engagement.

### GlnR-TAC is stabilized by complex protein-protein interactions between GlnR and the conserved domains of RNAP

In addition to the GlnR-DNA interactions described above, GlnR-TAC is also stabilized by distinctive and complex protein-protein interactions between GlnR and the conserved domains of RNAP. In *M. tuberculosis* GlnR-TAC, GlnR^I^_DBD not only recognizes the b2 site located downstream of the −35 element, but also interacts with the β flap domain from RNAP holoenzyme (**Figure 4B**), which has not been observed before. Residues R122, H163, R166, V167 and Q172 from GlnR^I^_DBD as well as L764, E768, G810, E811 and E831 from the loop structure of β flap mediated the interactions between GlnR^I^_DBD and β flap through salt bridges, hydrogen bonds, and van der Waals forces, which may act as a glue interface to stabilize the GlnR-TAC. Moreover, the structure also reveals additional interactions between GlnR^II^_DBD and σ^A^R4 of RNAP. Residues R187, T186, G184, D181 and H175 from GlnR^II^_DBD form six potential hydrogen bonds and salt bridges with residues E501, Q505, S508, K509 and S512 from the helix of σ^A^R4, and this interface is also stabilized by the hydrophobic and van der Waals forces contributed by residues F183, G184, A171 from GlnR^II^_DBD (**Figure 4C**). In comparison with σ^A^R4 alone in *M. tuberculosis* RPo (PDB ID: 6VVY), this interface between GlnR^II^_DBD and σ^A^R4 partially occludes σ^A^R4 from binding to the −35 element (**Supplementary Figure S8**). This well coincides with the previous observations from the crystal structure of ternary complexes including PhoB_DBD, σR4, β-flap-tip-helix and DNA (PDB ID: 3T72) (54), and suggests a common charge-based code between transcription factors and σ^A^R4. In accordance with this, substitutions of the key residues involved greatly reduced GlnR-dependent transcription activity and formation of GlnR-TAC as demonstrated by *in vitro* transcription assays and EMSA experiments, especially to residues H163 and R166 (**Figure 3F; Supplementary Figure S9B and S9C**). Additionally, though increasing evidences establish that αCTD plays an important role in bacterial transcription activation, αCTD is mostly invisible in transcription activation complexes due to its high flexibility, especially at the upstream DNA far away from the −35 element (29,32). Nevertheless, αCTD of *M. tuberculosis* RNAP is visualized in GlnR-TAC adjacent to the upstream promoter DNA, contacting the region covering a1 site. Residues D285, R259 and N288 from αCTD form three hydrogen bonds with the DNA backbone phosphate (**Figure S10A, S10B**). This distinctive αCTD-DNA interface provide a new proof for the highly versatile transcription activation modes of RNAP αCTD in bacterial transcription regulation, which functions directly with the upstream DNA instead of with activators or with UP-element DNA by the conserved 265 determinant in the previous reported transcription activation complexes, such as the classical CAP-TAC and the versatile SoxS-TAC (27–29,34). In addition, it seems that the electron densities in the map could not accommodate four GlnR_RECs, no matter what conformations they adopted. We tried to dock more GlnR_RECs into the densities based on the previous reported crystal structures of GlnR_REC, KdpE-DNA and PmrA-DNA (20,55,56). Fortunately, we finally succeeded in fitting GlnR^I^_REC and GlnR^IV^_REC into the densities (**Supplementary Figure S10C**). Intriguingly, a possible hydrophilic interaction interface is constituted and stabilized by residues T98, E102, and R106 from GlnR^IV^_REC and residues R61, R57 and D87 from GlnR^I^_REC through salt bridges and hydrogen bonds (**Supplementary Figure S10C, S10D**). Meanwhile, it is highly worth noting that GlnR^I^_REC may partially contact αNTD^I^, with a burying surface area of ~80 Å^2^ (**Supplementary Figure S10E**). Residues L21, S20, D13, and Y11 from GlnR^I^_REC and residues R55, Y96, E137, E158, and R161 from αNTD^I^ forms this first-reported interface through salt bridges, hydrogen bonds and Van der Waals forces, which might serve as a bridge to connect GlnR^IV^_DBD to the αNTD from RNAP core enzyme (**Supplementary Figure S10F**). Consistent with the aforementioned possible interactions, mutations of the key residues involved above apparently decrease transcriptional activities in the Mango-based transcription assays and formation of GlnR^I^_REC, confirming their essential roles in facilitating GlnR-dependent transcription activation (**Supplementary Figure S9; Supplementary Figure S10G**).

### Four GlnR molecules kinetically activate formation of a competent GlnR-TAC

To verify the kinetic stoichiometry of GlnR in *M. tuberculosis* GlnR-TAC, we measured the number of GlnR molecules involved via single-molecule fluorescence assay. To perform these measurements, we prepared a mutant of GlnR with a relocated and surface-accessible cysteine (GlnR C60SL147C, designated as GlnR147) and fluorescently labeled it with Alexa Fluor 647 C2-maleimide (GlnR147-Alexa647) (see Methods for details). DNA molecules labeled with Cy3 at +2 position of the non-template strand was assembled with *M. tuberculosis* RNAP and GlnR147-Alexa647 and then subjected to single-molecule fluorescence assays (**Figure 5A**). An alternative laser excitation (ALEX) module was implemented to detect the fluorescence of Cy3-DNA and GlnR147-Alexa647 respectively (**Figure 5B**). To qualify the number of GlnR147-Alexa647 in the complex, we increase the red laser (640 nm laser) power to 30 mW to induce quick photobleaching of Alexa647 dye and the number of steps that Alexa647 bleached would reflect the number of GlnR147 in the complex. Finally, we observed 139 out of 149 photobleaching events displaying one step and 10 other events displaying two steps (**Figure 5C**). This result gives a mean number of steps of 1.07 ± 0.02 (standard error of mean, SEM). Taking into account the dye-labeling stoichiometry of 27%, the stoichiometry of GlnR147-Alexa647 involved in *M. tuberculosis* GlnR-TAC is 4.0 ± 0.1 (SEM). This is in good agreement with our observations of the cryo-EM structure of GlnR-TAC, and also with the *in vitro* transcription assays which displayed a no better transcription activity for *amtB6* DNA containing 6 GlnR binding sites than for *amtB4* DNA just consisting of the downstream 4 GlnR binding sites **(Supplementary Figure S3C**). All of these suggest that four GlnR molecules kinetically and preferentially activates formation and stabilization of a more competent GlnR-TAC.

## Discussion

Whether bacteria can survive, grow or colonize in the various changing environments depends on their fine-tuned adaptive responses. Two-component systems (TCSs) composed of histidine kinases (HKs) and response regulator (RRs), are regarded as one of the most elaborate and efficient way to successfully fulfil stress adaptation at the transcription level (9,10,12). TCSs are ubiquitous in archaea, bacteria, lower eukaryotic organisms and plants but absent in mammals, therefore, bacterial TCSs are promising targets for drug discovery and design (57). The OmpR/PhoB subfamily transcription factors in bacteria are regarded as typical RRs involved in primary, secondary metabolisms, and have been extensively studied for four decades. However, most of these investigations focused on genetic characterization of their targeted promoter regulons, or on biochemical and crystal structural studies of the OmpR/PhoB subfamily RRs alone (DrrB, DrrD, RegX3, PrrA, MtrA, et al) (58–62) or in complex with their cognate *cis* element DNA (PhoB_DBD-DNA, full length KdpE and DNA, full length PmrA and DNA, et al) (52,53,56), a whole clear blueprint of how the OmpR/PhoB subfamily RRs activate transcription by coordinately engaging promoter DNA and RNAP holoenzyme is still lacking. By combination of our biochemical and structural data of GlnR-TAC, it is conceivable for us to get in-depth understanding of the above long-standing question. On one hand, four GlnR protomers engage promoter DNA in GlnR-TAC mainly through interactions between GlnR_DBD and the corresponding *cis* element DNA, along with retaining interactions between two adjacent GlnR_DBDs, as being identified in the co-crystal structure of *sae*GlnR_DBD in complex with its *cis* element DNA. This distinctive “head-to-tail” mode of DNA engagement not only demonstrates and expands our understanding on promoter recognition of the reported transcription factors, but also provide a general model for the OmpR/PhoB subfamily response regulators. This is in good agreement with the high binding affinity between GlnR and DNA identified in the EMSA experiment **(Supplementary Figure S9**). On the other hand, complex protein-protein interactions between different GlnR molecules protomers and the conserved domains of RNAP efficiently facilitate transcription activity and formation of GlnR-TAC. GlnR^I^_DBD and GlnR^II^_DBD interact with the β flap and σ^A^R4, respectively, which are consistent with the previous reported transcription factors (54). It is intriguing that αCTD also contacts the upstream a1 site DNA, and αNTD interacts with GlnR^IV^_DBD which were further verified by our mutational analyses. These novel interfaces **(Supplementary Figure S10**) provide new evidences for the global and versatile regulation modes of the highly conserved domains of RNAP (especially the whole α subunit) in bacterial transcription initiation, by acting as specialized platforms to interact with various transcription factors. Notably, this distinctive architecture simultaneously possesses the general features of the reported canonical Class I and Class II activators, with the upstream two GlnR protomers finely contact α subunit and promoter DNA mainly in the way as Class I activators (28,34), while the downstream two GlnR protomers coordinately and deftly engages its cis-element DNA, and the conserved β flap, σ^A^R4 of RNAP as Class II activators (27,29,31,33). Thus, the biochemical and structural data provide new insights into the underlying transcription activation mechanisms for the OmpR/PhoB subfamily proteins, and reveal a unique mode of bacterial transcription regulation.

Although the GlnR *cis*-elements having been studied for many years, by bioinformatics analyses and experimental studies, the GlnR binding consensus sequences seem to be more complex and diverse, especially on the number and importance of different GlnR binding sites. A 22-bp GlnR binding consensus element, including four sites (a1, b1, a2, b2 sites) separated by 6 bases, was first proposed for GlnR *cis*-element on the basis of a dozen studies (21). Then Wang *et al*. amended this typical GlnR binding element and put forward only three of the four GlnR binding sites seems to be essential for GlnR binding to DNA *in vitro* and to regulate the gene transcription *in vivo*, while a2 site is unnecessary (22,23). In line with these, various kinds of strong interactions between GlnR-DBDs and the typical GlnR binding consensus sites were extensively verified in our structure of GlnR-TAC. From our cryo-EM structure, we could get the following possible explanations for the dispensability of a2 site: the a2 site is not only engaged by GlnR^II^_DBD, but also retained by the σ^A^R4 conserved domain. When the a2 site is mutated, the σ^A^R4 domain may still recognize the non-optimal −35 element DNA, and strengthen the stabilities of GlnR-TAC in combination with the upstream GlnR-DNA interactions. Another positive regulatory role for the a2 site might be recruiting GlnR to DNA consensus sites ahead of RNAP engagement.

As to the kinetic activation mechanism of GlnR, the previous biochemical investigation of *A. mediterranei* GlnR proposed that GlnR might activate transcription via two conformations—complex I is formed by GlnR binding to both the a1 and b1 sites of the GlnR dependent operon, and it subsequently promotes formation of a transcription competent complex II by engaging the other GlnR binding sites (23). This regulation mode is quite similar to that of the LysR-type transcription regulators (LTTRs), such as LysR, catR and CbbR. These regulators firstly form complex I by a dimer binding to the regulatory binding site (RBS), then the other dimer binds to the ABS (activation binding site) and retains in a tetramer state through their N terminal dimerization domains, and finally assemble into an active complex II to facilitate transcription (23,63). In support of the above mentioned, both the cryo-EM structure and single-molecule FRET assays of GlnR-TAC showed that four GlnR protomers structurally and kinetically promote formation of a more stable and competent GlnR-TAC, reflecting a predominant active conformation for GlnR proteins. This is consistent with our unsuccessful trials in obtaining another cryo-EM structure of GlnR-TAC containing six GlnR binding sites (a3-b3, a1-b1 and a2-b2 sites), which also showed the same four GlnR protomers bound to the downstream a1-b1 and a2-b2 sites of *amtB* promoter (data not shown), indicating of lower binding affinity of GlnR to the upstream a3-b3 sites than to the downstream ones. This could also be explained by the competitive binding activities between GlnR and PhoP (a global regulator for phosphate metabolism) uncovered by Wang *et al* (64), which may play physiological roles in coordinating and responding to both signals of nitrogen and phosphate availability in bacteria. Taking all these into consideration, it is most probably that GlnR-dependent transcription activation is established as a novel and elaborate transcription activation mechanism through co-regulation of the “head-to-tail” assembled GlnR tetramer upon environmental stimuli (**Figure 5D**), the GlnR tetramer may be the most stable form for the physiological function of GlnR. Existence of the extra a3-b3 site may not be predominantly regulated by GlnR, which may act as an additional transcription regulation strategy to promptly respond to environmental nitrogen availability, and orchestrately shares precise cross-talk with the essential phosphate metabolism. However, this needs to be further explored in the future, both *in vitro* and *in vivo*.

## Data Availability

Accession number for cryo-EM density map: EMD-34816 for GlnR-TAC (Electron Microscopy Data Bank). Accession number for atomic coordinates: 8HIH for GlnR-TAC,8HML for *Sae*GlnR complexed with DNA (Protein Data Bank).

## Supplementary Data

Supplementary Data are available at PNAS Online.

## Acknowledgements

We appreciate Shenghai Chang at the Center of Cryo-Electron Microscopy in Zhejiang University School of Medicine and Guangyi Li, Fangfang Wang, Liangliang Kong at National Center for Protein Science, Shanghai for assistance with cryo-EM sample preparation and data collection. We thank the Core Facilities, Zhejiang University School of Medicine for technical support. We thank the Experiment Center for Science and Technology, Nanjing University of Chinese Medicine for experimental assistance.

This work was funded by the National Natural Science Foundation of China (82072240, 81903756, 32270192, 32270037, 32000025), Jiangsu Province of China (BK20211302 to J.S.), the Open Project of Chinese Materia Medica First-Class Discipline of Nanjing University of Chinese Medicine (No. 2020YLXK008 to W.L., No. 2020YLXK016 to J.S.), the Fok Ying Tung Education Foundation, Open Funding Project of the State Key Laboratory of Bioreactor Engineering and Jiangsu Specially-Appointed Professor Talent Program to W.L. This work was also funded is supported by the National Natural Science Foundation of China (No. 12004420, 32071228), the Strategic Priority Research Program of the Chinese Academy of Sciences (No. XDB37000000), and the Youth Innovation Promotion Association of CAS (No. 2021009).

## Author Contributions

J. S., ZZ. F., JC. X, FF. L., YQ. Z., A.J. W., FL. W., Q.S, and L. W. performed the experiments. A.J. W., J. S. performed cryo-EM sample preparations and data collections. F. Y., W. L. performed cryo-EM structure determination. J. S., W. L. designed the study, W. L., J. S., Y. F., and S. W. analyzed data, and wrote the paper.

