## Supplementary information for "Structural insights into transcription activation mechanism of the global regulator GlnR from actinobacteria"

**Running title:** GlnR-dependent transcription activation complexes.

>***amtB6* DNA**

GCACGACGACGACCGGCCCCGGCCGTTCACCCACGCGTAACACGCACCGTGCCTT
CGTCACGGCGGCGAAACAACGAGGGGCTTCCACCGAAACCGCGCTGCGTCAATG
TCGTGGCGC**A**TAACCGGCCCACCCCTCACCGCCCGCCCGCCAGGCACGTACGAAG
GAAGGATTGGTATGTGGTATATTCGTACGTGCC

>***amtB4* DNA**

ACCGTGCCTTCGTCACGGCGGCGAAACAACGAGGGGCTTCCACCGAAACCGCGC
TGCGTCAATGTCGTGGCGC**A**TAACCGGCCCACCCCTCACCGCCCGCCCGCCAGGC
ACGTACGAAGGAAGGATTGGTATGTGGTATATTCGTACGTGCC

>***narG* DNA**

CCGTCGCTGTTAGGAAACCGACGGTGTGGTTGACGGTGGCCGCCGTCAACTTGGT
TAGAACAACGTGACAAAACGTTAACTTGGGTTTGCATGCCCGTAGCGATTACGATG
GTTTT**C**TGGACGCGTGGCGACAACTTCCGGGCAGGGGCACGTACGAAGGAAGGA
TTGGTATGTGGTATATTCGTACGTGCC

GlnR binding box　–35 element　–10 element　**transcription start site**

Transcription product　mango

**Figure S1. The sequences of promoter DNA used.**

The GlnR binding box, –10 element, transcription product, *mango* sequence are highlighted in yellow, purple, grey, and cyan, respectively. The –35 element is underlined. The transcription start site is shown in red and bold font.

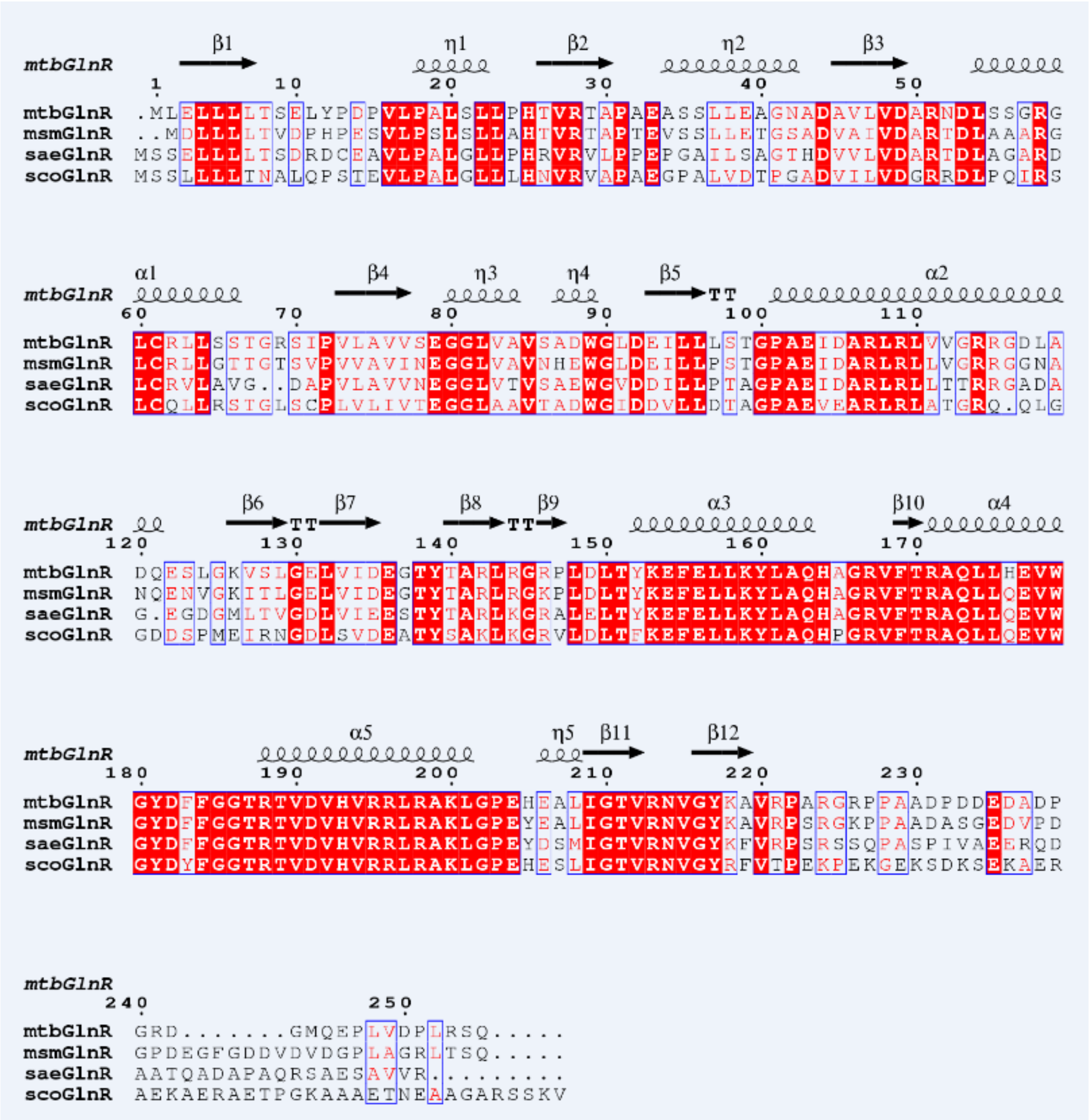


**Figure S2. Structural based sequence alignment of GlnR and its orthologues**.

Amino acid sequences of *M. tuberculosis* GlnR (mtbGlnR), *Mycobacterium smegamatis* GlnR (*msm*GlnR), *Saccharopolyspora erythraea* GlnR (*sae*GlnR), and *Streptomyces coelicolor GlnR* (*sco*GlnR) were aligned. The invariant residues among them are highlighted in red, and conserved amino acids are boxed. The secondary structure elements of mtbGlnR are shown at the top and as the template in the ESpript Web server (1).

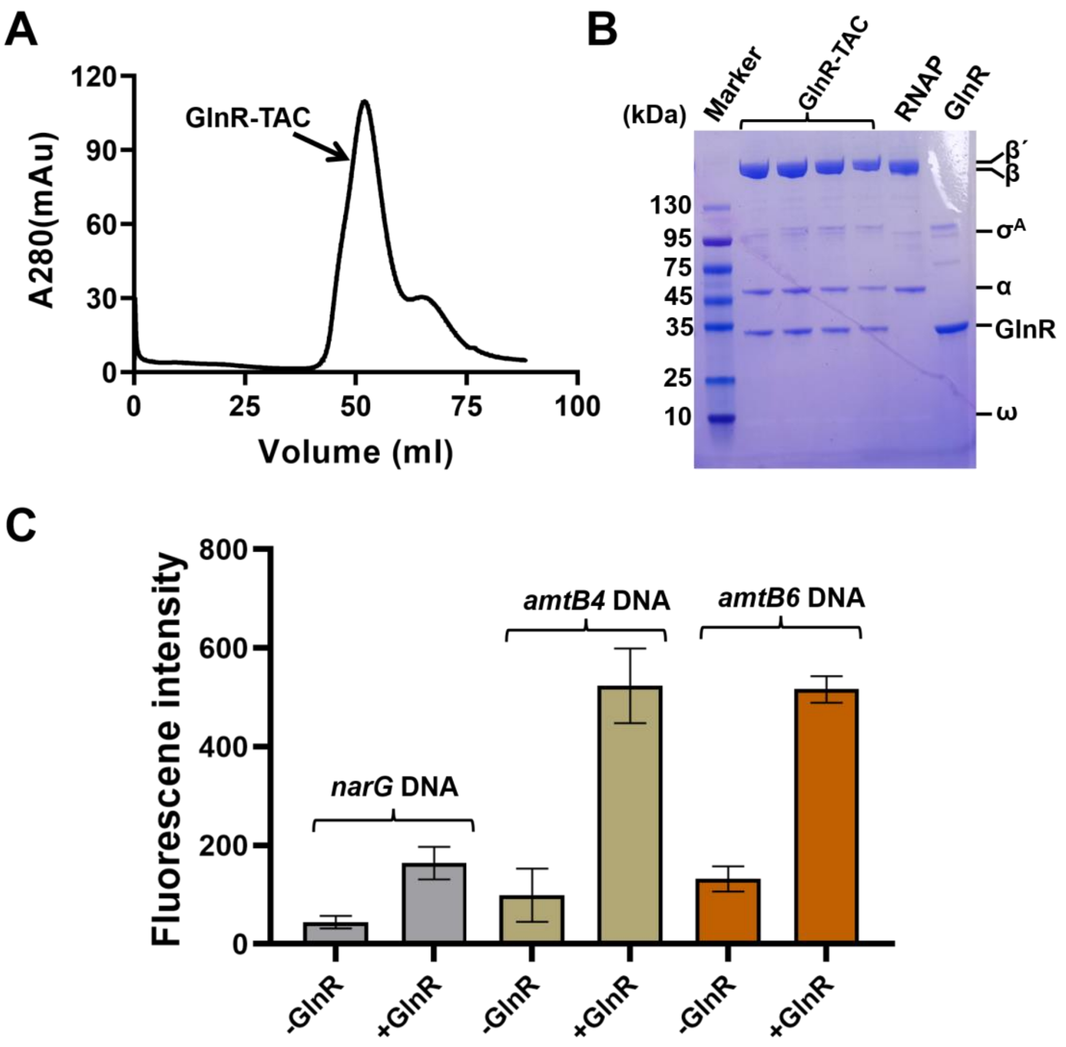


**Figure S3. Purification and verification of *M. tuberculosis* GlnR-TAC.**

**(A)** Gel filtration map of *M. tuberculosis* GlnR-TAC.

**(B)** SDS-PAGE of the purified GlnR-TAC complex.

**(C)** Relative transcription activity of *M. tuberculosis* GlnR dependent transcription activation complex determined by *in vitro* transcription assay on different DNA. 0.1 μM *M. tuberculosis* RNAP, 30 nM mango-ended DNA and 16 μM *M. tuberculosis* GlnR were included in the reaction except others indicated in *Materials and Methods*. Data for *in vitro* transcription assays are means of 3 technical replicates. Error bars represent mean ± SEM of n = 3 experiments.

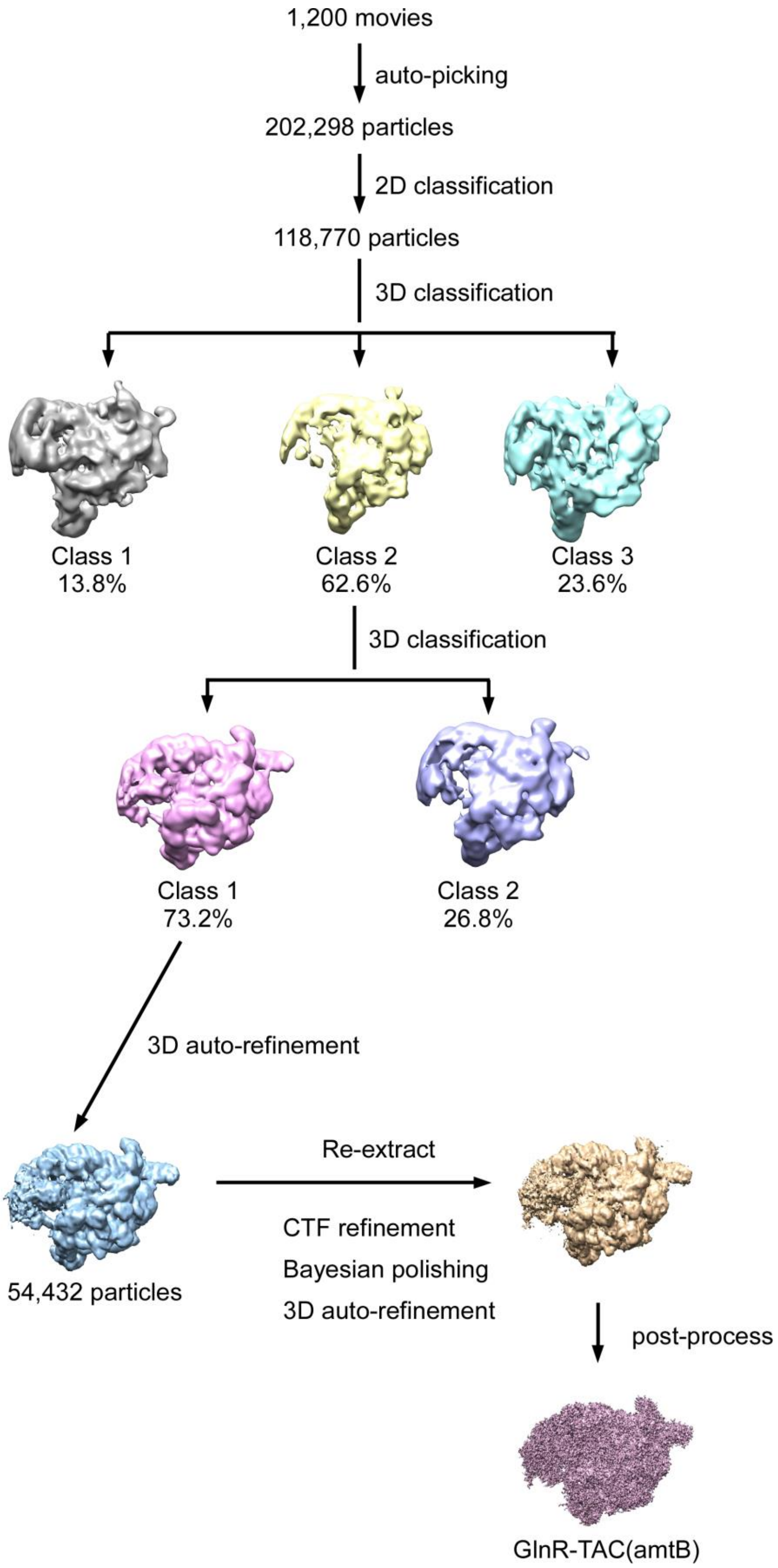


**Figure S4. Data processing pipeline for the dataset of *M. tuberculosis* GlnR-TAC.**

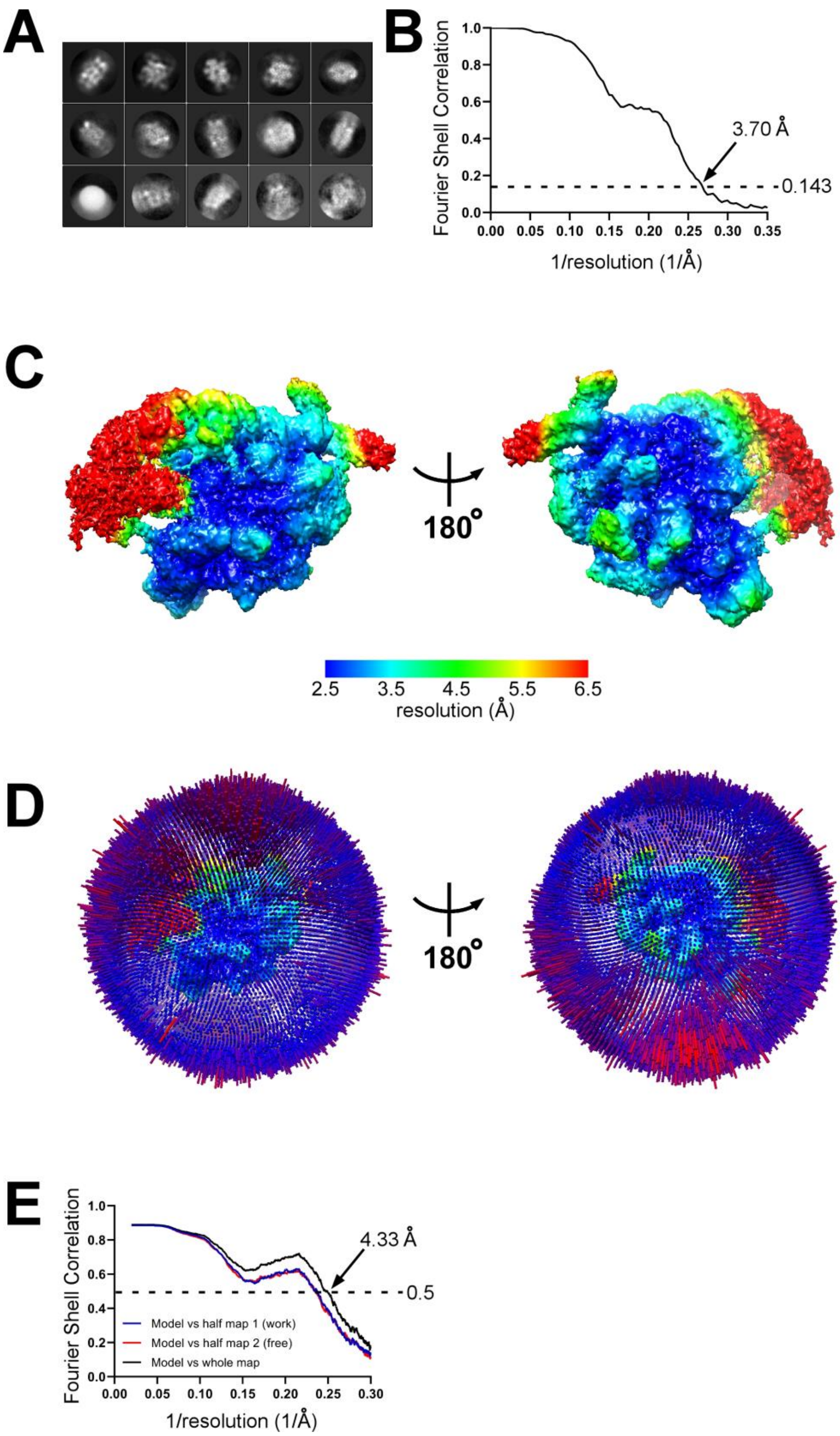


**Figure S5. Cryo-EM data of *M. tuberculosis* GlnR-TAC.**

**(A)** Representative classes from 2D classification.

**(B)** Gold-standard FSC. The gold-standard FSC was calculated by comparing the two independently determined half-maps from RELION. The dashed line represents the 0.143 FSC cutoff (2), which indicates a nominal resolution of 3.70 Å.

**(C)** Cryo-EM density map colored by local resolution. Local resolution calculation was performed using blocres (3). View orientation as in **Figure 2B**.

**(D)** Angular distribution of particle projections. View orientations as in **(C)**.

**(E)** FSC calculated between the model and the half map used for refinement (work), the other half map (free), and the full map.

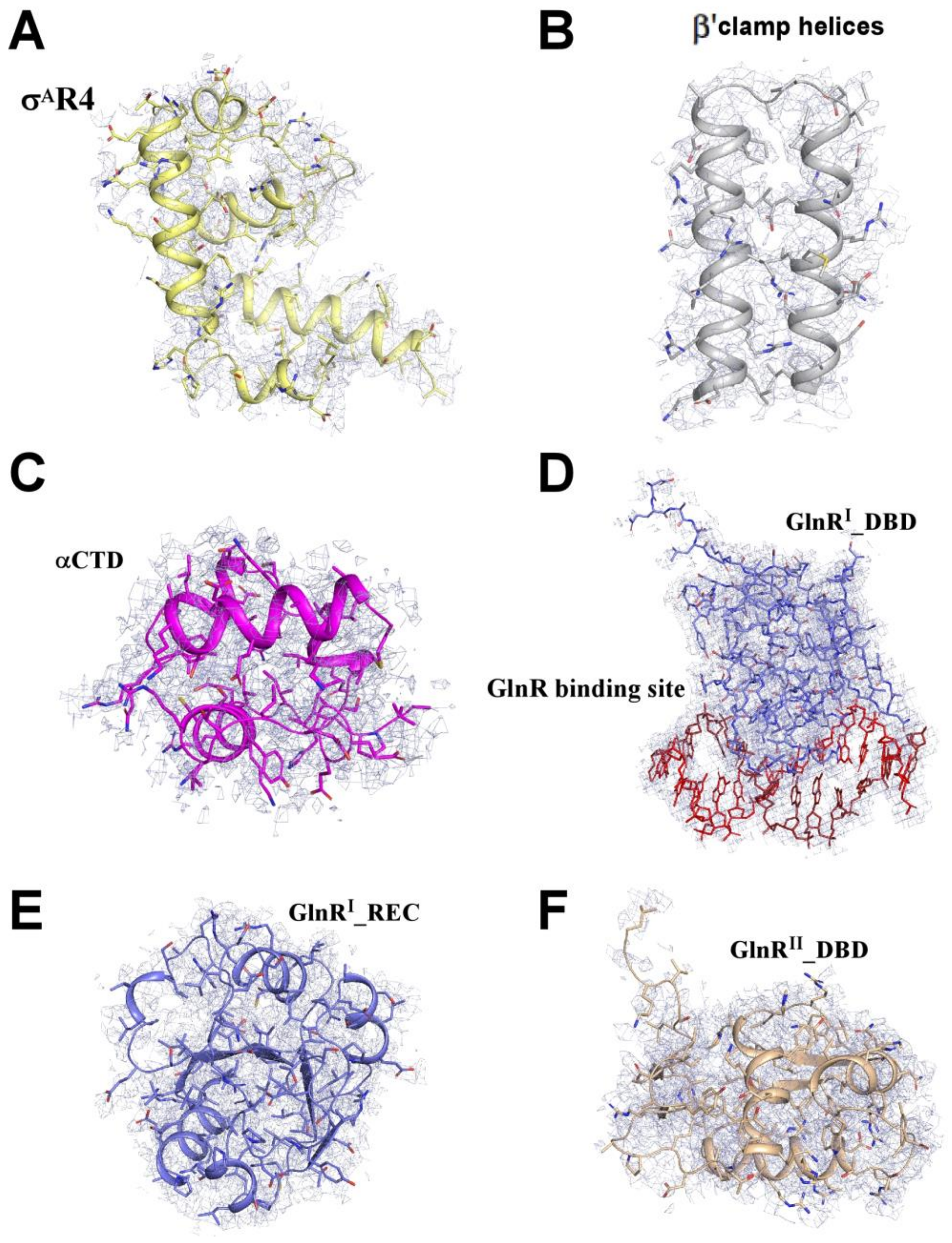


**Figure S6. Representative cryo-EM densities of superimposed models in *M. tuberculosis* GlnR-TAC.**

(A) Cryo-EM density map (grey mesh) and the superimposed model (yellow cartoon) of $\sigma^A$R4. **(B)** Cryo-EM density map (grey mesh) and the superimposed model of β′ clamp helices. **(C)** Cryo-EM density map (grey mesh) and the superimposed model of αCTD. **(D)** Cryo-EM density map (grey mesh) and the superimposed model of $\text{GlnR}^{I}$_DBD and its DNA binding site. **(E)** Cryo-EM density map (grey mesh) and the superimposed model of $\text{GlnR}^{I}$_REC. **(F)** Cryo-EM density map (grey mesh) and the superimposed model of $\text{GlnR}^{II}$_DBD. Other colors are shown as in **Figure 2**.

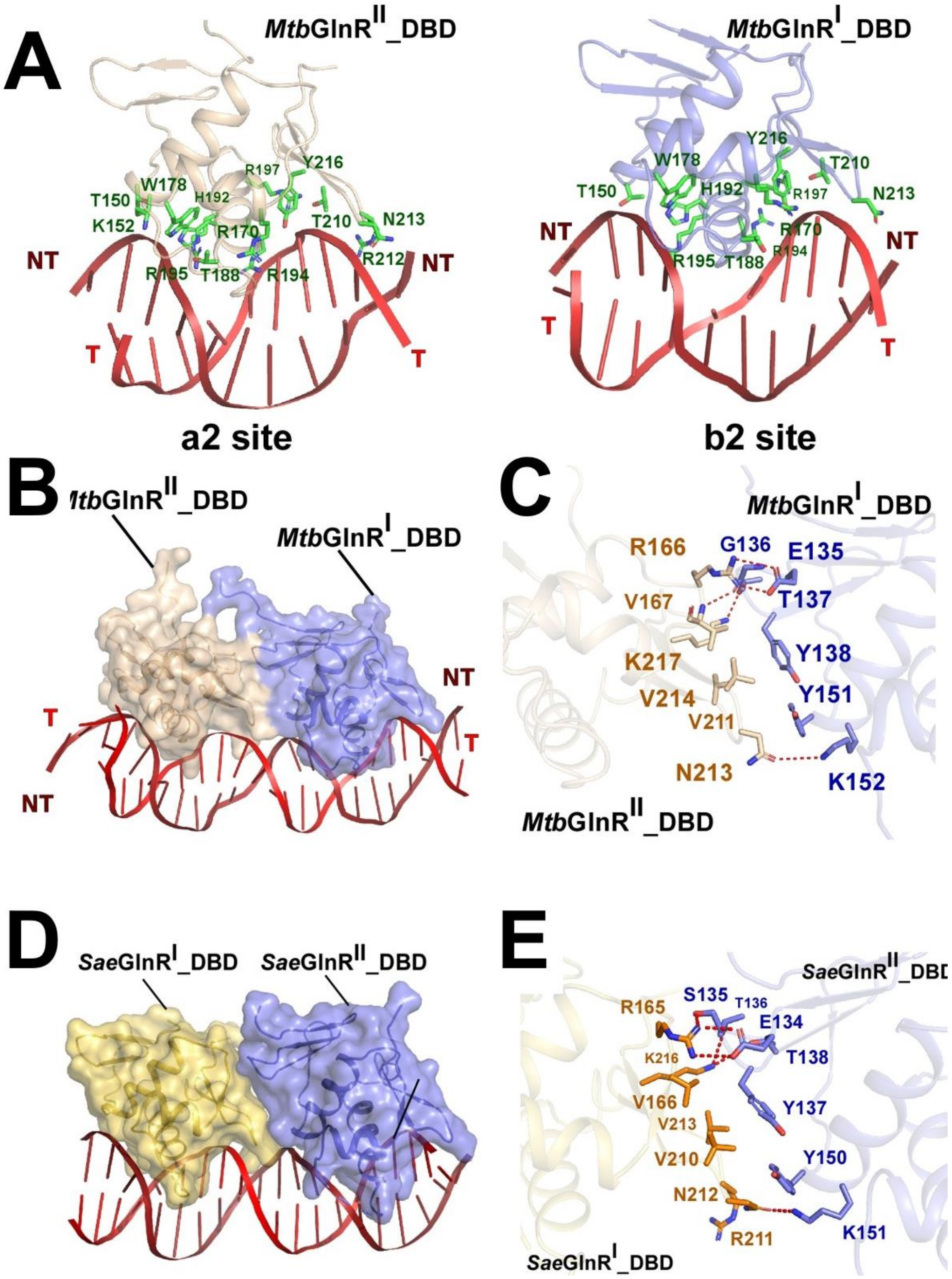


**Figure S7. The protein-DNA and protein-protein interactions between two adjacent GlnR_DBDs and the GlnR binding sites in GlnR-TAC.**

**(A)** Relative locations of GlnR$^{II}$_DBD, GlnR$^{I}$_DBD and their corresponding upstream double-stranded GlnR binding sites a2 and b2. **(B-C)** The relative locations and detailed interactions between GlnR$^{I}$_DBD and GlnR$^{II}$_DBD bound to the promoter DNA. **(D-E)** The relative locations and detailed interactions between *Sae*GlnR$^{I}$_DBD and *Sae*GlnR$^{II}$_DBD bound to the promoter DNA. Polar interactions are shown as red dashed lines. Colors are shown as in **Figure 2**.

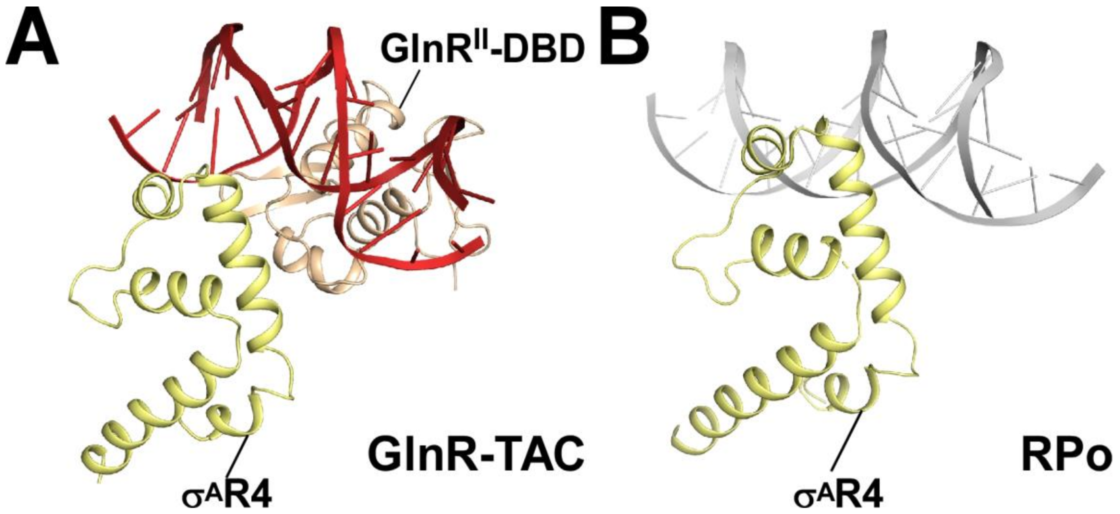


**Figure S8. The interactions between GlnR, σ$^{A}$R4 and -35 element DNA.**

**(A)** Relative locations of GlnR$^{II}$_DBD, σ$^{A}$R4, and the upstream double-stranded DNA in *M. tuberculosis* GlnR-TAC. **(B)** Relative locations of σ$^{A}$R4 and the upstream typical -35 element DNA in *M. tuberculosis* RPo (PDB ID: 6VVY) (4).

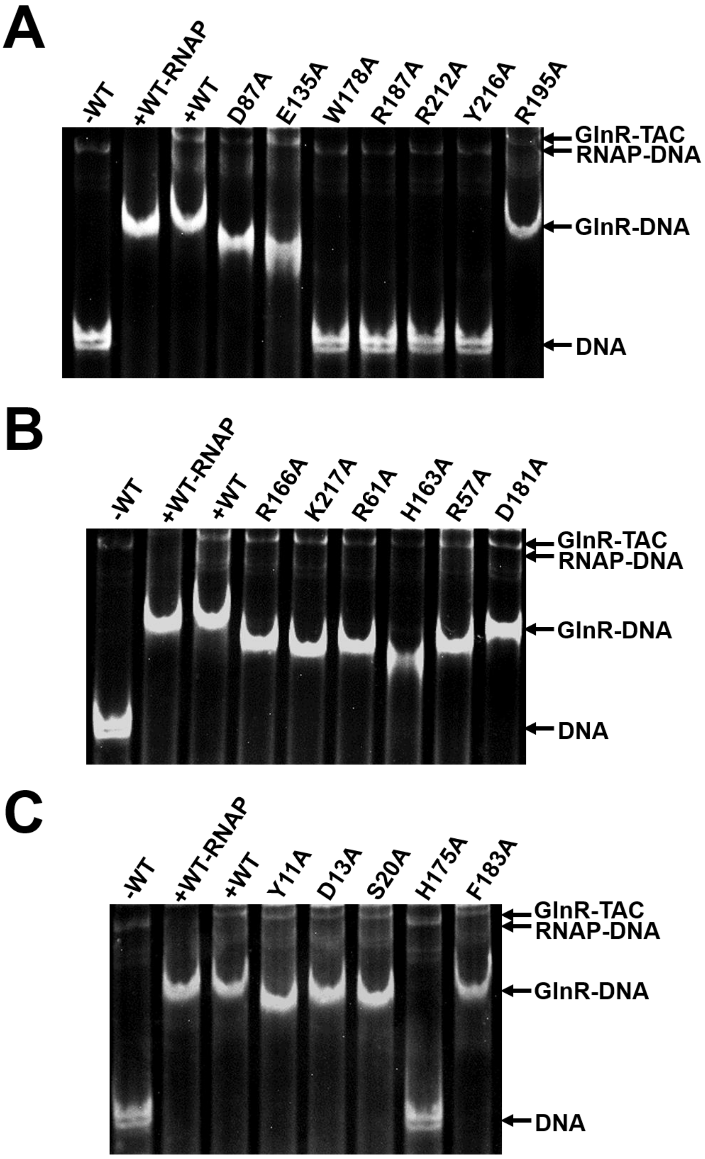


**Figure S9. GlnR derivatives show defects on formation of GlnR-TAC.**

**(A)** Electrophoretic mobility shift assay for wild-type GlnR and its mutants mainly involved in GlnR-DNA and GlnR-GlnR interfaces. **(B)** Electrophoretic mobility shift assay for wild-type GlnR and its mutants mainly involved in GlnR-β flap and GlnR-$\sigma^A$R4 interfaces. **(C)** Electrophoretic mobility shift assay for wild-type GlnR and its mutants mainly involved in GlnR-GlnR, GlnR-αCTD and GlnR-$\sigma^A$R4 interfaces. Reaction conditions are described in detail in *Materials and Methods*. Bands of GlnR-TAC, RNAP-DNA, GlnR-DNA and free DNA are showed on the right, respectively.

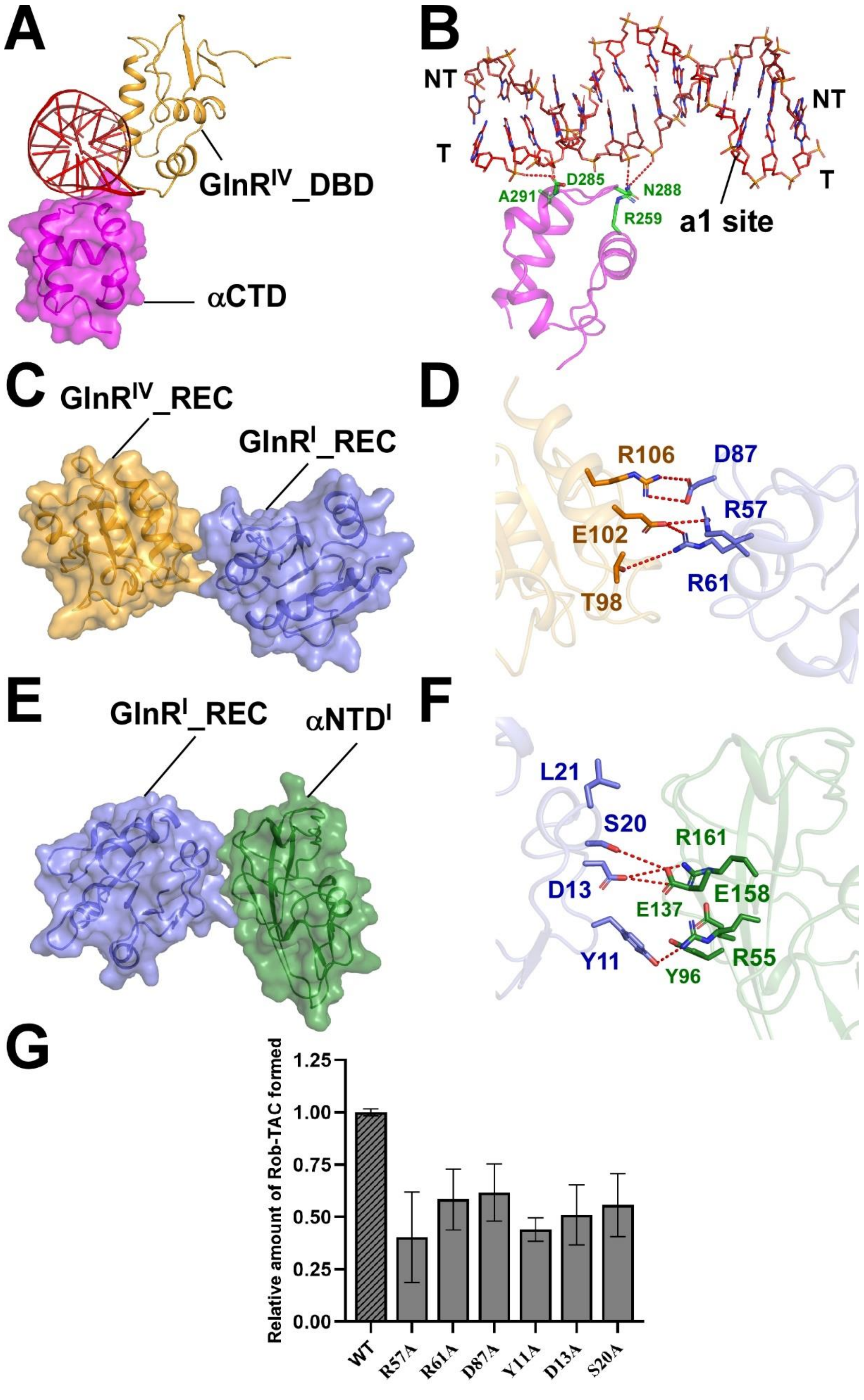


**Figure S10. The possible critical interactions between *M. tuberculosis* GlnR$^{IV}$,**

**GlnR$^{I}$, αCTD and DNA.**

**(A)** Relative locations of *M. tuberculosis* GlnR$^{IV}$_DBD, RNAP αCTD and the upstream double-stranded DNA of GlnR binding site a1. **(B)** Detailed interactions between *M. tuberculosis* RNAP αCTD and the double-stranded DNA of GlnR binding site a1. **(C-D)** Relative locations and detailed interactions of *M. tuberculosis* GlnR$^{IV}$-REC and GlnR$^{I}$-REC. **(E-F)** Relative locations and detailed interactions of *M. tuberculosis* GlnR$^{I}$-REC and αNTD$^{I}$. **(G)** Mutational effects of the above key GlnR residues identified by *in vitro* transcription assays. Reaction conditions are described in detail in *Materials and Methods*. Colors as in **Figure 2**.

### Supplementary Materials and Methods

#### Fluorescent labeling of GlnR147

The purified GlnR147 and Alexa Fluor 647 C2-maleimide (Invitrogen) were incubated at a molar ratio of 1:5 in labeling buffer (10 mM Tris-HCl pH 8.0, 75 mM NaCl and 5 mM $MgCl_2$) at a final volume of 110 µl. This reaction was performed at 4 ℃ for 12 hr protecting from light and stopped by adding DTT at a final concentration of 1 mM. Free dyes were separated using size-exclusion chromatography (HiLoad superdex200, GE healthcare) in labeling buffer containing 5% (v/v) glycerol and 1 mM DTT. Protein concentration and dye-labeling stoichiometry were determined using nanodrop with extinction coefficients of 19940 $M^{-1}cm^{-1}$ for GlnR147 and 265000 $M^{-1}cm^{-1}$ for Alexa647 dye. Labeling stoichiometry was 27%. Labeled GlnR147 was aliquoted and flash frozen in liquid nitrogen and stored at -80 ℃.

#### DNA construct preparation

HPLC purified DNA oligos (amtBpNT+2Cy3 and amtBpT, the underlined "T" is labeled with Cy3) for single-molecule fluorescent experiments were purchased from Sangon Biotech. These two DNA oligos were annealed at 1:1 ratio by heating to 95 ℃ for 5 min followed by slowly cooling down to room temperature in transcription buffer (40 mM Tris-HCl pH 8.0, 100 mM NaCl and 10 mM $MgCl_2$ and 1 mM DTT). A final concentration of 0.5 mg/ml BSA was then added. The annealed DNA construct was aliquoted and stored at -20 ℃. Sequences of amtB+2Cy3 oligos are shown in **Supplementary Table S1**.

#### Preparation of DNA-RNAP-GlnR complex

This complex was prepared by mixing Mtb RNAP with DNA construct at final concentrations of 110 nM and 56 nM respectively in a final volume of 10 µl. The sample was incubated at 37 ℃ for 30 min with shaking (550 rpm). GlnR-Alexa647 was then added at a final concentration of 1.8 µM in a final volume of 50 µl. The sample was then incubated at 37 ℃ for 30 min with shaking (550 rpm) which can be subjected to single-molecule fluorescence measurements.

**Supplementary Table S1. Primer sequences used in this study.**

| **Primer name** | **Sequence (5′ to 3′)** |
|---|---|
| *amtB6*_F1 | GCACGACGACGACCGGCCCCGGCCGTTCACCCACGCGTAACAC |
| *amtB6*_F2 | GTTCACCCACGCGTAACACGCACCGTGCCTTCGTCACGGCGGCGAAACAACGAGGGGC |
| *amtB6*_F3 | GGCGAAACAACGAGGGGCTTCCACCGAAACCGCGCTGCGTCAATGTCGTGGCGCATAAC |
| *amtB6*_F4 | CAATGTCGTGGCGCATAACCGGCCCACCCCTCACCGCCCGCCCGCCAGGCACGTAC |
| *amtB4*_F | ACCGTGCCTTCGTCACGGC |
| *amtB*_R | CACATACCAATCCTTCCTTCGTACGTGCCTGGCGGGCGG |
| *mango*_R | GGCACGTACGAATATACCACATACCAATCCTTCCTTC |
| *narG*_F0 | CCGTCGCTGTTAGGAAACCGACGGTGTGGTTGACGGTGGCCGCCGTCAACTTGGTTAGA |
| *narG*_F1 | CCGCCGTCAACTTGGTTAGAACAACGTGACAAAACGTTAACTTGGGTTTGCATGCCC |
| *narG*_F2 | TAACTTGGGTTTGCATGCCCGTAGCGATTACGATGGTTTTCTGGACGCGTGGCGACAAC |
| *narG*_F3 | CTGGACGCGTGGCGACAACTTCCGGGCAGGGGCACGTACGAAGGAAGGATTGGTATGTG |
| *amtB* scaffold_T | TGCATCCGTGAGTCGAGGGTAATAAACGCAGCGCGGTTTCGGTGGAAGCCCCTCGTTGTTTCGCCGCCGTGACGAAG |
| *amtB* scaffold_NT | CTTCGTCACGGCGGCGAAACAACGAGGGGCTTCCACCGAAACCGCGCTGCGTTATAATGGGAGCTGTCACGGATGCA |
| *amtB*pNT+2Cy3 | GCCCCGGCCGTTCACCCACGCGTAACACGCACCGTGCCTTCGTCACGGCGGCGAAACAACGAGGGGCTTCCACCGAAACCGCGCTGCGTCAATGTCGTGGCGCA**T**AACCGGCCCACCCCT |
| *amtB*pT | AGGGGTGGGCCGGTTATGCGCCACGACATTGACGCAGCGCGGTTTCGGTGGAAGCCCCTCGTTGTTTCGCCGCCGTGACGAAGGCACGGTGCGTGTTACGCGTGGGTGAACGGCCGGGGC |
